# Simultaneous multifunctional transcriptome engineering by CRISPR RNA scaffold

**DOI:** 10.1101/2022.06.21.497089

**Authors:** Zukai Liu, Paul Robson, Albert Cheng

## Abstract

RNA processing and metabolism are subjected to precise regulation in the cell to ensure integrity and functions of RNA. Though targeted RNA engineering has become feasible with the discovery and engineering of the CRISPR-Cas13 system, simultaneous modulation of different RNA processing steps remains unavailable. In addition, off-target events resulting from effectors fused with dCas13 limit its application. Here we developed a novel platform, <u>*C*</u>ombinatorial <u>*R*</u>NA <u>*E*</u>ngineering via <u>*S*</u>caffold <u>*T*</u>agged gRNA (CREST), which can simultaneously execute multiple RNA modulation functions on different RNA targets. In CREST, RNA scaffolds are appended to the 3’ end of Cas13 gRNA and their cognate RNA binding proteins are fused with enzymatic domains for manipulation. We show that CREST is capable of simultaneously manipulating RNA alternative splicing and A-to-G or C-to-U base editing. Furthermore, by fusing two split fragments of the deaminase domain of ADAR2 to dCas13 and PUFc respectively, we reconstituted its enzyme activity at target sites. This split design can reduce more than 90% of off-target events otherwise induced by a full-length effector. The flexibility of the CREST framework will enrich the transcriptome engineering toolbox for the study of RNA biology and the development of RNA-focused therapeutics.

## INTRODUCTION

Post-transcriptional regulation controls gene expression at the RNA level, and its dysfunction is involved in many diseases(1). Leveraging hundreds of RNA binding proteins, a cell regulates an RNA transcript in various biological processes, including its maturation, modification, stability, and localization to ensure proper function. Failure of any of these steps might result in cellular dysfunction. For example, RNA alternative splicing, occurring in as many as 80% of genes, contributes the most to transcriptomic diversity(2). Another example of post-transcriptional regulation is in adenosine-to-inosine RNA editing, which not only affects translation by altering codons, but also changes the recognition of splice sites(3). Therefore, the study and manipulation of the coordination between different RNA processing steps are keys to understanding the complicated network of post-transcriptional regulation and treating RNA malfunction related diseases.

A variety of tools based on RNA scaffolds were developed for transcriptome engineering in the last two decades. An RNA scaffold is an RNA motif with a specific sequence or structure which can be recognized by a given RNA binding domain (RBD). Several pairs of RNA scaffolds and their RBDs were well-characterized, including Pumilio, MS2 and PP7 systems(4). Fusions of these RBDs with functional effectors have been used for the study of RNA biology, including RNA location, live imaging, and RNA translational regulation (5–12). However, these RNA scaffold-based technologies have several key limitations. For example, the MS2 system requires the integration of MS2 sequence into the transcripts of interest by genetic engineering, which is time and labor intensive. In addition, the insertion of these RNA scaffolds may alter the dynamics and functions of the target transcripts(13–15). Although PUF can be engineered to recognize different octamers, the diversity of this is limited(10, 16).

Recent discoveries of novel CRISPR-Cas13 systems overcome the limitations of RNA scaffold-based tools and enable efficient and precise targeting of endogenous RNAs(17–19). The CRISPR-Cas13 system consists of two components, a guide RNA (gRNA) with sequence complementary to the target transcript and the Cas13 protein with endonuclease activity. Importantly, a catalytically inactive Cas13 mutant (dCas13) can be fused with different regulators and enzymes for targeted RNA manipulation with high specificity(20–29). However, current CRISPR-Cas13-based tools are limited to one specific function for one dCas13-effector in the same cell. Although dual-color imaging of different transcripts with two orthogonal dCas13 proteins was reported, the number of characterized Cas13 orthologs is limited and their large sizes increase the challenge of co-delivering multiple dCas13 proteins into the same cell(21). These hamper the development of multifunctional RNA engineering tools.

An additional challenge in RNA manipulation is the poor substrate selectivity of the effectors. For example, adenosine deaminase acting on RNA (ADAR) binds to double-stranded RNA and then converts adenosine (A) to inosine (I) via deamination. The resultant inosine is functionally equivalent to guanosine (G)(30). Fusion of the deaminase domain of ADAR2 (ADAR2-DD) with dCas13 can achieve site-specific A-to-G editing. However, dCas13-ADAR2-DD showed substantial dCas13-independent transcriptome-wide off-target activity(17). The reconstitution of a split enzyme at a given locus provides the opportunity to limit off-target events elicited intrinsically by the full-length enzyme(31). With Cas13-split ADAR2-DD direct fusions, this goal can be accomplished by two nearby gRNAs but RNA secondary structures on target transcripts can potentially inhibit enzyme reconstitution.

Here, we report a novel platform called <u>*C*</u>ombinatorial <u>*R*</u>NA <u>*E*</u>ngineering via <u>*S*</u>caffold <u>*T*</u>agged gRNA (CREST) by combining RNA scaffold with CRISPR-Cas13. As proof-of-principle, we engineered orthogonal MCP- and PUF-based CREST modules for splicing modulation. We then created CREST base editing modules for A-to-G and C-to-U editing. We further generated high-efficient split RNA editing modules using CREST architecture and showed the significant reduction of off-target events in the split version compared with full-length effector. Finally, we demonstrated that orthogonal CREST modules could be used for multifunctional transcriptome engineering, specifically manipulating simultaneous alternative splicing and base editing. The CREST platform will enable complex operations within RNA regulatory networks relevant to both fundamental research and clinical applications.

## RESULTS

### Design of CREST

CREST comprises gRNA appended with RNA scaffolds, nuclease- and gRNA-processing-deficient ddPspCas13b (hereafter as ddCas13b) and RNA binding domain (RBD) fused with effector. The 5’ of the gRNA (spacer) is designed to be complementary to the target transcript while its 3’ end is appended with RNA scaffolds, acting as the bridge between ddCas13b and RBD-effector fusions (Supplementary Figure S1B). To achieve multifunctional transcriptome modulation, we fused different effectors with specific RBDs and switched the scaffold in the gRNA to control the engineering consequences.

### CREST-mediated RNA alternative splicing modulation

We first tested CREST architecture on the modulation of alternative splicing. *SMN2* was selected as the target because the inclusion of exon7 in *SMN2* is a well-recognized treatment strategy for Spinal Muscular Atrophy. We recently reported the induction of exon 7 inclusion in *SMN2* mRNA by dCas13b-splicing factor direct fusion(20). As previously reported, we used three gRNAs designed to target downstream of exon 7 of the pCI-SMN2 minigene to induce exon 7 inclusion (Figure 1A). To adopt the CREST architecture, the three gRNAs were tagged with different numbers of MS2 or PBSc scaffolds that can recruit splicing effectors built by replacing RNA binding motif of RBFOX1 with MCP or PUFc, respectively. We co-transfected the pCI-SMN2 minigene with dCas13b, gRNA-MS2/MCP-RBFOX1 or gRNA-PBSc/PUFc-RBFOX1 into HEK293T cells and analysed splicing activity using quantitative RT-PCR. Initially, we failed to achieve splicing activation (Figure 1B, 1C, grey columns). We reasoned that the intrinsic gRNA processing activity of (d)Cas13b required for releasing individual mature gRNAs from polycistronic pre-gRNA may cleave away the MS2 and PBS scaffolds, and the binding of dCas13 without effectors to *SMN2* minigene transcript showed minor inhibitory effect on exon 7 inclusion. Since the amino acids required for PspCas13b gRNA processing was unknown, we aligned PspCas13b with PbuCas13b in which K399 was reported as required for gRNA processing(32).

**Figure 1.**
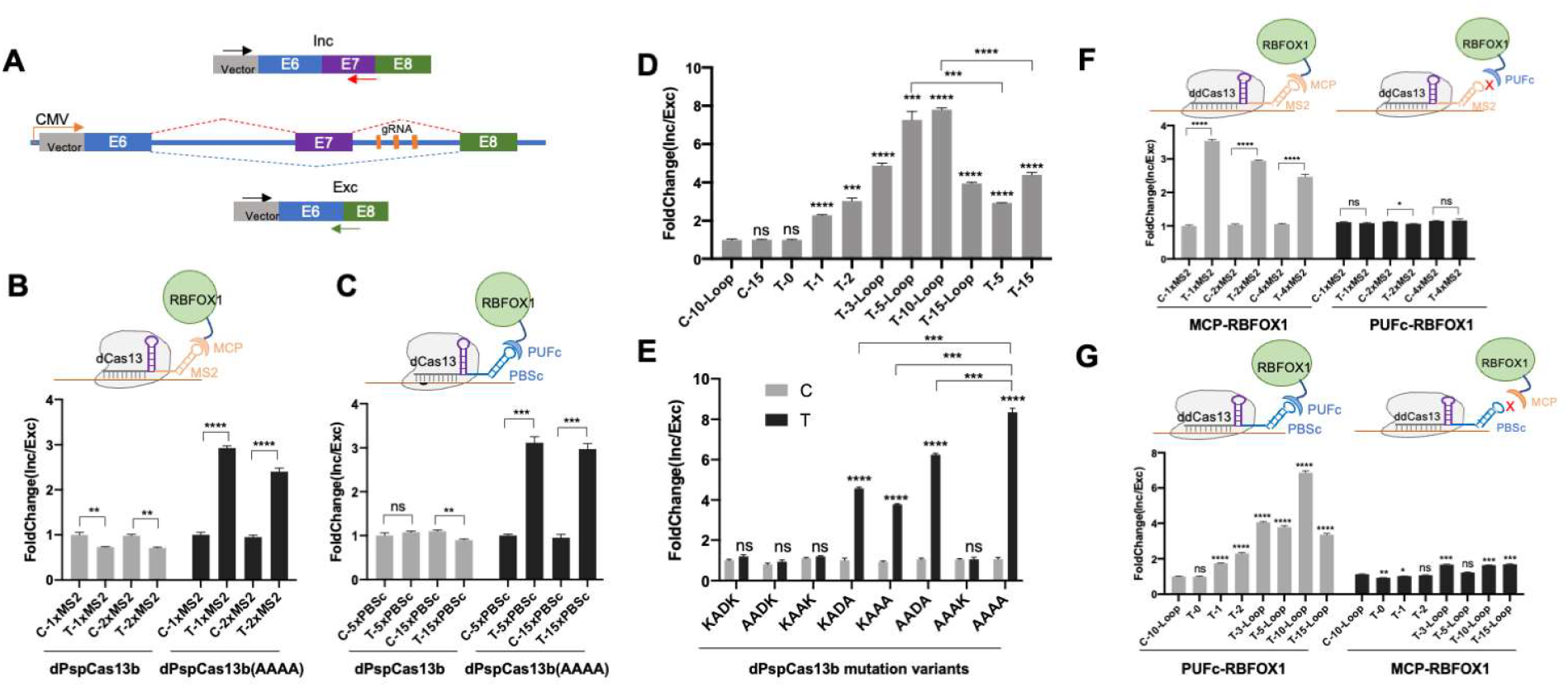
CREST-mediated RNA alternative splicing modulation. (A) Schematic of pCl-SMN2 minigene reporter containing the genomic region of spanning *SMN2* exon 6 to exon 8 downstream of a CMV promoter. Inclusion and exclusion isoforms can be quantified with the same forward primer annealing to a constitutive region (Black arrow) and a reverse primer annealing to isoform-specific exon junctions (red arrow for the inclusion isoform and green arrow for the exclusion isoform). Three gRNAs targeting downstream intron of exon 7 are marked in orange. (B) Above: Schematic of CREST MCP-MS2 system. MS2 scaffold was tagged at the 3’ end of gRNA and MCP was fused with RBFOX1 without RRM domain for induction of exon7 inclusion. Bottom: splicing readouts measured by RT-qPCR. Y axis: Exon inclusion efficiency (inclusion/exclusion ratio measured by PCR) is represented as fold change relative to non-target control gRNA. X axis: HEK293T cells were co-transfected with pCl-SMN2 reporter and different dpspCas13 mutants, gRNAs and different number of MS2 scaffold. Nontarget control (C-) and minigene target -specific (T-) gRNA are indicated. The copies (1× or 2×) of the MS2 scaffold are indicated. Different dPspCas13b variants were showed as indicated. dPspCas13b: deficiency at targeting transcripts cleavage. dPspCas13b(AAAA): deficiency at both targeting transcripts cleavage and gRNA processing. (C) Above: Schematic of CREST PUFc-PBSc system. PUFc binding sites (PBSc) were appended to the 3’ end of gRNA and PUFc was fused with RBFOX1 without RRM domain. Bottom: splicing readout measured by RT-qPCR. 5 and 15 copies of PBSc with ‘GCC’ linker were tested. (D) Optimization of PBSc linkers. Fold change of Inclusion and Exclusion ratio was normalized by non-target control gRNA with 10 copies of PBSc with loops (C-10-Loop). ‘Loop’ stands for the high GC content stem-loop structure between PBSc and the copy number (0-15) of the PBSc motifs are indicated. PBSc with indicated number but without ‘Loop’ were linked by ‘GCC’. dPspCas13b(AAAA) was used in all groups. (E) Screen for the minimal set of mutations in dPspCas13b compatible with CREST. The original motif (KADK) for gRNA processing in pspCas13b was used as control and other variants were tested as indicated. Gray (C) stands for non-target control gRNA and black (T) stands for on-target gRNA. PUFc-RBFOX1 and 10 copies of PBSc with Loop were used in all conditions. dPspCas13b with AAAA mutation was referred as ddCas13 thereafter. (F) Test of the crosstalk between the PUFc and MS2. HEK293T cell were co-transfected with ddCas13, pCI-SMN2 minigene, gRNA-MS2 and MCP-RBFOX1 (Gray columns on the left) or PUFc-RBFOX1(Black columns on the right) as indicated. Y axis indicated the fold change of Inclusion and Exclusion ratio relative to non-target control. (G) Test of the crosstalk between the MCP and PBSc. HEK293T cell were co-transfected with ddCas13, pCI-SMN2 minigene, gRNA-PBSc and MCP-RBFOX1(Gray columns on the left) or PUFc-RBFOX1(Black columns on the right) as indicated. All data were displayed as mean ± S.E.M, n = 3. *P<0.05, ** P < 0.01,***P<0.001, ****P<0.0001, ns, not significant, by two-sided t-test.

A stretch of four amino acids (KADK) containing three charged residues K367, D369 and K370 on PspCas13b aligned around K399 of PbuCas13b and were thus selected as candidates for PspCas13b gRNA processing activity. We therefore mutated all charged residues into the non-charged amnio acid alanine, changing KADK to AAAA (Supplementary Figure S1C). dCas13b with the AAAA mutation was able to induce the inclusion of exon 7 with the CREST architecture (Figure 1B and 1C). To our surprise, the number of RNA scaffolds had little effect on the inclusion efficiency. We suspected multiple scaffolds on the gRNA may not fold independently, thus reducing the number of scaffolds available for RBD-effector binding. We thus set out to optimize gRNA design. RNAfold predicted that 5 and 15 copies of PBSc interspaced with a GCC linker formed unwanted secondary structures which may interfere with PUFc binding (Supplementary Figure S2E&F)(33). We replaced the GCC linker between PBSc with high GC content stem loops to free PBSc sites from secondary structure in the low energy state (Supplementary Figure S2A-D). The result showed that 5 and 10 copies of PBSc with stem loops performed the best (Figure 1D). We next investigated the minimal set of mutations in dCas13b within the KADK stretch required for CREST architecture by screening all the possible combinations of mutation in KADK. The dCas13b variant with an AAAK mutation was unable to induce inclusion of exon 7, indicating that K370 is essential and sufficient for gRNA processing. While dCas13b with K370A mutation allowed splicing activity in the context of CREST, the additional K367A and D369A mutations improved exon 7 inclusion efficiency (Figure 1E). Therefore, we used the dCas13b variant with AAAA mutation in our CREST architecture and referred it as ddCas13b to signify both target cleavage and gRNA processing deficiency.

The orthogonality between different RNA scaffolds is key to multifunctional RNA engineering. Using alternative splicing as a readout, we evaluated orthogonality of CREST by co-transfecting all pairwise combinations of gRNA with different RNA scaffolds and RBD-RBFOX1. We showed that MCP-RBFOX1 but not PUFc-RBFOX1 activated the inclusion of exon 7 in the presence of gRNA-MS2 (Figure 1F). Similarly, gRNAs tagged by PBSc were unable to induce exon 7 inclusion to any meaningful degrees in the presence of MCP-RBFOX2 (Figure 1G). Taken together, these results demonstrated not only that CREST can be engineered to control alternative splicing in mammalian cells, but also that orthogonality of different RNA scaffold systems form the basis of CREST-mediated multifunctional RNA engineering.

### CREST-mediated A-to-G base editing

Programmable RNA base editing shows clinical significance to disease treatment by correcting gene mutations at the RNA level. We chose RNA editing as the second function to evaluate CREST. To provide a convenient readout of RNA editing activity, we generated a reporter with the CMV promoter-driven Clover and mutant mScarlet transgenes separated by a T2A peptide(34). The mutation in mScarlet encodes a premature stop codon (UAG), marking transfected cells with green fluorescence only if the transcript remains unedited. The test editing event changes the UAG codon to UGG, effectively rescuing mScarlet expression and giving cells dual fluorescence (Figure 2A, Supplementary Figure S3). With this reporter, A-to-G editing efficiency can be assessed by flow cytometry. We tagged gRNAs with PBSc so that when complexed with ddCas13b, these gRNAs would recruit PUFc-ADAR2-DD to the target RNA. Given that ADARs selectively deaminate adenosines of A-C mismatches in the context of double-stranded RNA, we placed a cytidine in the gRNA at the position facing the target adenosine on the target transcript(35). A previous study revealed that the length of gRNA and mismatch distance (first matched nucleotide to the editing site) are critical to the editing efficiency(17). We used gRNAs with a 30nt spacer and tiled gRNA positions for mismatch distances from 2nt to 30nt. Consistent with the previous report, we found that 22-26 nt were the optimal mismatch distance, which showed nearly 60% editing efficiency (Figure 2B). To examine whether the editing resulted from the recruitment of endogenous ADARs by gRNAs independent of exogenous effectors as reported in LEAPER(36), we transfected cells with gRNAs in the absence of PUFc-ADAR2-DD and did not observe any editing events (Figure 2B). Though A-to-G conversion mostly happens in the cytoplasm, the addition of nuclear export signal (NES) to ddCas13b and PUFc-ADAR2-DD did not improve editing efficiency, indicating the “baseline” localization of these proteins is sufficient to support editing activity (Figure 2B).

**Figure 2.**
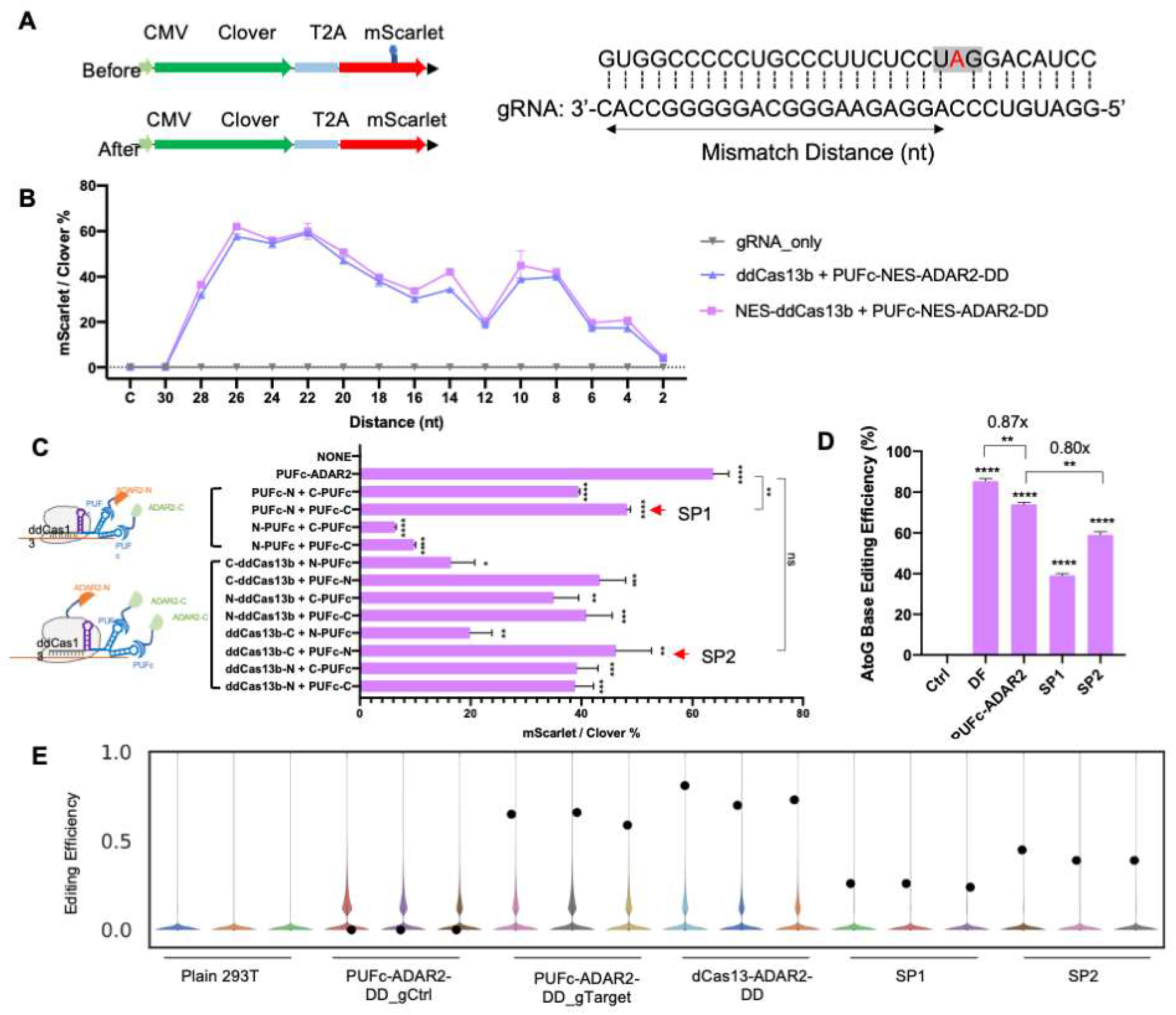
CREST-mediated A-to-G base editing. (A) Left: schematic of the reporter system for A-to-G editing. CMV driven Clover and mScarlet ORFs were separated by a T2A cleavage site. Before editing, the premature stop codon (UAG) disrupts the complete translation of mScarlet and made the transfected cell Clover-positive only (top). A-to-G editing converted UAG to UGG to rescue the production of mScarlet and turned cell double positive for Clover and mScarlet (bottom). Right: target sequence within the mutant mScarlet (top sequence) and spacer sequence in gRNA (bottom sequence). Mismatch distance is defined as the space between first matched nucleotide to the target adenine as indicated by double arrow. (B) Tilling of mismatch distances from 2nt to 30nt in gRNAs for A-to-G editing. HEK293T cells were co-transfected with reporter, gRNA with or without ddCas13, PUFc-ADAR2-DD as indicated. Flow cytometry was done 48 hours after transfection and the editing efficiency was calculated by the ratio of mScarlet and Clover positive cells (Y axis). X-axis: the mismatch distance of spacer sequence in gRNAs, C stands for non-target control gRNA. Different transfection groups are shown as indicated at the right. All gRNAs were tagged by 10xPBSc-Loop. (C) Reconstitution of split ADAR2-DD by CREST system. ADAR2-DD was split into two fragments (N terminal and C terminal) and fused with PUFc (Top left) or ddCas13 and PUFc separately (Bottom left). X-axis: Editing efficiency measured by flow cytometry. Y-axis: different orientations and combinations of fusion proteins were indicated. NONE: cell transfected with gRNA only was used as negative control. The intact ADAR2-DD fused with PUFc (PUFc-ADAR2) was used as a positive control. N stands for the N-terminal fragment of ADAR2-DD and C stands for the C-terminal fragment of ADAR2-DD. All gRNAs were tagged by 3×PBSc-Loop with 22nt mismatch distance. (D) A-to-G editing efficiency quantified by sanger sequencing. Ctrl: HEK293T cell transfected with reporter, ddCas13, PUF-ADAR2-DD and non-target control. DF: direct fusion of ADAR2-DD with original dCas13b without AAAA mutation and on-target gRNA without RNA scaffolds. PUFc-ADAR2: HEK293T cell transfected with reporter, ddCas13, PUF-ADAR2-DD and on-target gRNA tagged by 3×PBSc-Loop. SP1 and SP2 are indicated in (C). (E) Violin plots representing distribution of A-to-G editing yields observed at reference sites. Plain HEK293T cell without transfection was used as control. Effectors transfected into HEK293T cells were as indicated and black dots indicate editing yields at the target site within the mScarlet transcript. Three replicates were done for each group. Data were displayed as mean ± S.E.M, n = 3. *P<0.05, ** P < 0.01, ***P<0.001, ****P<0.0001, ns, not significant, by two-sided t-test.

We next tested if CREST architecture can be used to support split ADAR2-DD to reduce dCas13-independent off-target RNA editing events. We split ADAR2-DD into two parts, the N-terminal and C-terminal fragments as reported (37), and fused them with ddCas13b or PUFc. We speculated that CREST is able to reconstitute ADAR2-DD catalytical activity at the target locus in two scenarios. In the first scenario, PUFc is fused with either the N-or C-terminal fragments of ADAR2-DD and tandem PBSc on gRNA recruits multiple PUFc, which stochastically includes both N- or C-terminal fragments of ADAR2-DD for reconstitution. In the second scenario, different fragments of ADAR2-DD are fused with ddCas13b and PUFc respectively and gRNA functions as a bridge to bring them together. By screening all possible combinations, we found two optimal configurations, PUFc-N/PUFc-C and ddCas13b-N/PUFc-C, which were termed as SP1 and SP2 respectively (Figure 2C). As expected, the transfection of either split half ADAR2-DD was not sufficient for A-to-G editing (Supplementary Figure S4). To evaluate RNA editing activity among the different designs, we quantified editing efficiency by RT-PCR followed by Sanger sequencing(38). CREST-PUFc-ADAR2 showed comparable efficiency to the dCas13b-ADAR2 direct fusion and the CREST split version SP2 reaches almost 80% of the RNA editing efficiency achieved by CREST-PUFc-ADAR2(Figure 2D). Strikingly, deep RNA sequencing revealed significant reduction (90-97%) of off-target events in both SP1 and SP2 compared with CREST-PUFc-ADAR2 and dCas13-ADAR2 (Figure 2E). We identified more than 13,000 and 7,700 off-target events in cells transfected with CREST-PUFc-ADAR2 and dCas13-ADAR2, respectively, both having full-length ADAR2-DD but only 810 and 535 in the two CREST split-ADAR constructs, SP1 and SP2, respectively (Supplementary Figure S5).

In addition, we selected 5 genes with known disease-causing G-to-A point mutations to test the clinical potential of our CREST system. Some of these genes (*APC, MECP2* and *SMN1*) were reported to be corrected by direct fusion of dCas13 and ADAR2-DD at high efficiency but others (*CFTR* and *HBB*) were edited less than 10%(25). We used direct fusion to benchmark our CREST-PUFc-ADAR2-DD and SP2 and showed that both are robust in A-to-G editing for APC, *MECP2* and *SMN1*. Interestingly, the mutation in *CFTR* was only corrected by CREST-PUFc-ADAR2-DD at high efficiency (>20%) but not in direct fusion, indicating the advantage of CREST system in specific contexts (Supplementary Figure S6).

Taken together, we showed that our CREST system is able to mediate A-to-G base editing at high efficiency and provided evidence for decreased off-target effects through CREST-based split-and-reconstitution architecture.

### CREST-mediated C-to-U base editing

We subsequently applied CREST to C-to-U base editing. RESCUE-S, evolved from ADAR2-DD, induces C-to-U conversion in RNA duplexes but not A-to-G conversion(19). Therefore, we fused PUFc with RESCUE-S for C-to-U base editing. For readout, we chose a previously reported target sequence for further direct comparison(19), inserted it right after the mScarlet coding region in a CMV-mScarlet reporter, and measured the editing efficiency by Sanger sequencing (Figure 3A). Similar to what we observed for A-to-G editing, we found that gRNAs with 26nt mismatch distance showed the best efficiency (close to 80%, Figure 3B, Supplementary Figure S7). To address the concern that accumulated PUFc-RESCUE-S might increase bystander editing events, we assessed C-to-U conversion rate within the window of 30nt and found no significant changes in the adjacent cytosines (Figure 3C). In addition, we split and reconstituted the RESCUE-S enzyme by CREST. Given that the amino acids flanking the split site of ADAR2-DD are conserved in RESCUE-S, we adapted the same split site as used for our split ADAR2-DD above (Supplementary Figure S9). Unlike ADAR2-DD, only the hybrid combinations of split RESCUE-S worked with nearly 15% editing efficiency, including C-ddCas13b/PUFc-N, ddCas13b-N/C-PUFc and ddCas13b-N/PUFc-C (Figure 3D). This result indicates the conformation of RESCUE-S differs from ADAR2-DD and the split site should be optimized in the future studies. We tested CREST-mediated C-to-U base editing on three additional disease-relevant gene mutations and found that CREST-PUFc-RESCUE-S is able to induce Cytosine to Uridine conversion at comparable or higher efficiency as direct fusion of dCas13-RESCUE-S (Supplementary Figure S8).

**Figure 3.**
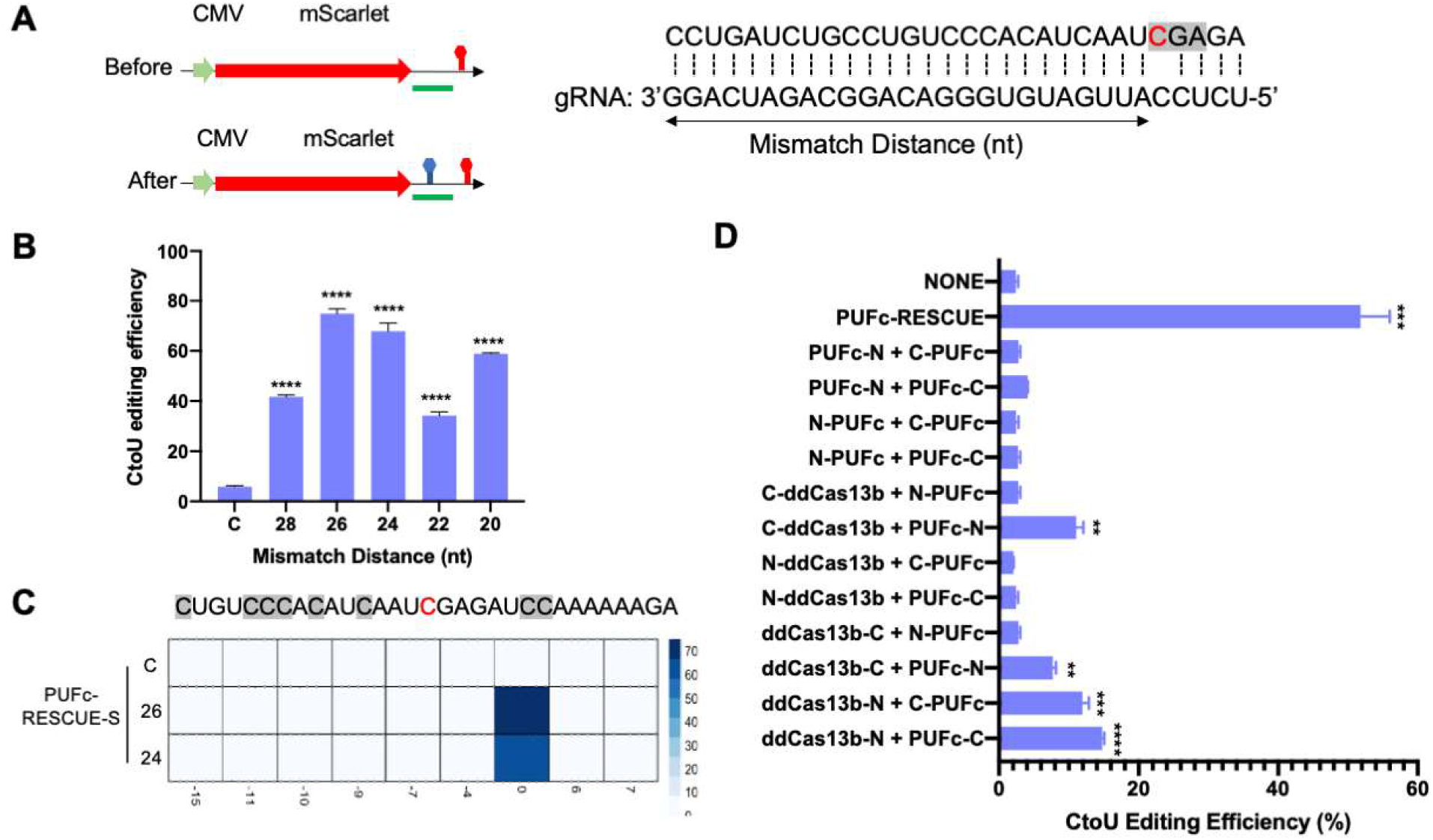
CREST-mediated C-to-U base editing. (A) Left: the diagram of reporter transgene for C-to-U editing. The target region was marked by a green bar. Right: target sequence (Top sequence) and spacer sequence in gRNA (bottom sequence). Mismatch distance is defined as indicated. (B) Tilling of mismatch distances from 20nt to 28nt in gRNAs for C-to-U editing. HEK293T cells were co-transfected by reporter transgene, ddCas13b, PUFc-RESCUE-S and different gRNAs with 10×PBSc-Loop. Y axis: C-to-U editing efficiency was quantified by RT-PCR followed by sanger sequencing. X axis: mismatch distances and C stands for non-target control gRNA. (C) Heatmap of C-to-U conversion rate of neighboring cytosines located within 15nt upstream and downstream of the target site. HEK293T cells were transfected with reporter, ddCas13, PUFc-RESCUE-S and gRNAs as indicated in each row. C stands for the control gRNA. The numbers 26 and 24 indicate mismatch distances. Editing efficiency was quantified by RT-PCR followed by sanger sequencing. (D) Reconstitution of split RESCUE-S by CREST. RESCUE-S was split into two fragments, C terminal and N terminal, then fused with PUFc only or ddCas13b and PUFc separately at different orientations as indicated. NONE: cell transfected with gRNA only was used as negative control. The intact RESCUE-S fused with PUFc (PUFc-RESCUE) was used as a positive control. N stands for the N-terminal fragment of RESCUE-S and C stands for the C-terminal fragment of RESCUE-S. All gRNAs were tagged by 10xPBSc-Loop. C-to-U editing efficiency was quantified by Sanger sequencing. Data were displayed as mean ± S.E.M, n = 3. *P<0.05, ** P < 0.01,***P<0.001, ****P<0.0001, ns, not significant, by two-sided t-test.

### CREST-mediated multifunctional RNA modulation

Complex RNA regulation in the cell involves not only the multiple processing and modification steps on the same transcript, but also the coordinated regulation of different transcripts. Comparing with Cas13-direct fusion effectors offering single functionality, we posit that CREST is able to perform multifunctional RNA modulation. To demonstrate this, we tested simultaneous alternative splicing and base editing as a proof-of-principle. The splicing and RNA editing reporters, ddCas13b and RBD-effectors were co-transfected into 293T cells, and the modulation outcome was controlled by the introduction of desired gRNAs with RBD-effector-matching scaffolds. We showed that the addition of gRNAs tagged by MS2 can only activate MCP-RBFOX1-mediated alternative splicing but not PUFc-ADAR2-mediated A-to-G editing and vice versa (Figure 4A, lane 2 and 3). Moreover, the addition of both gRNAs edited both reporters for two independent events at the same time at high efficiency (Figure. 4A, lane 4). Similarly, this orthogonal multifunctional engineering can be used for the alternative splicing and C-to-U base editing as well as the A-to-G and C-to-U base editing (Figure 4B&C). In sum, CREST enables us to perform multifunctional modulation and shows the potential to edit transcripts at different levels.

**Figure 4.**
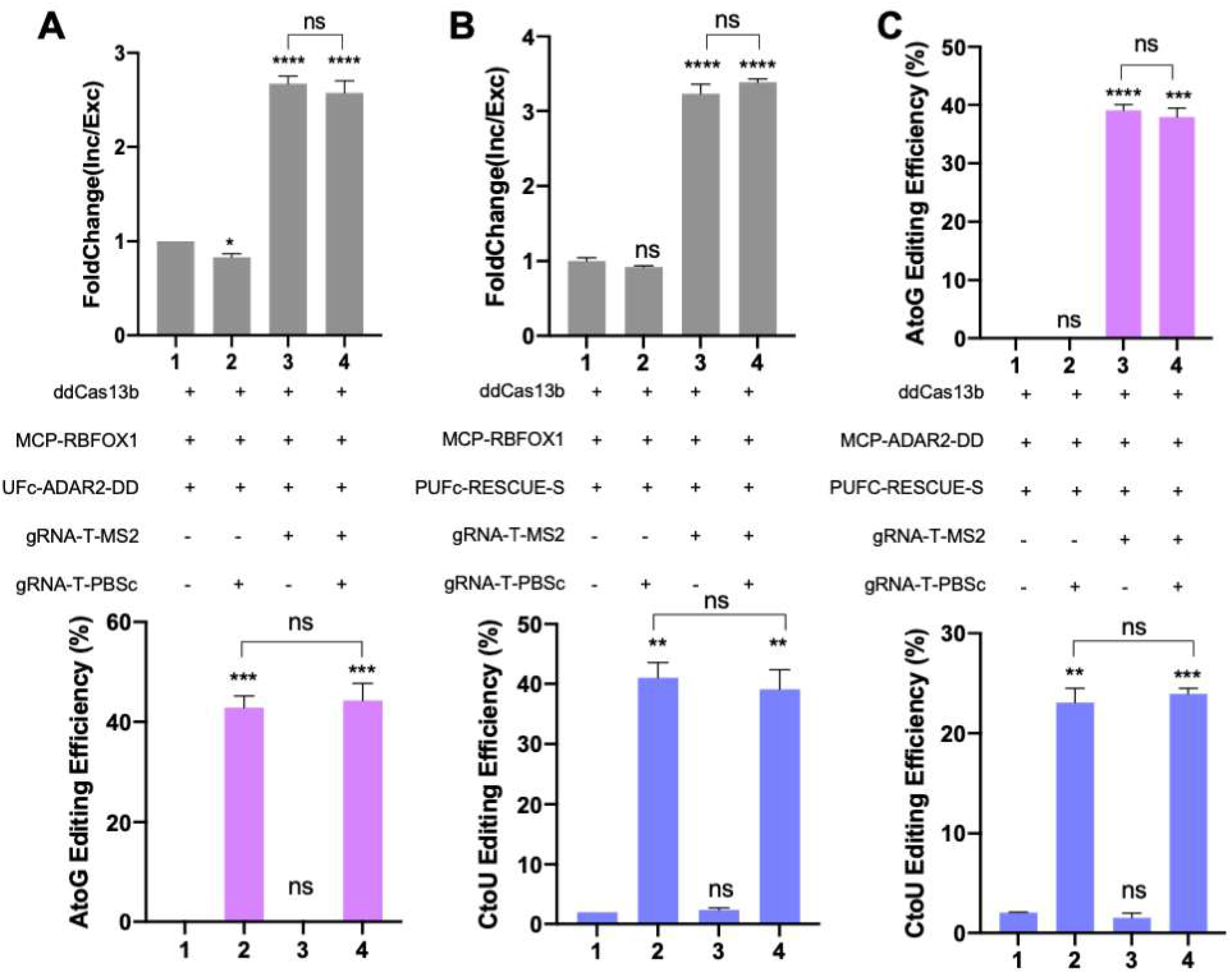
CREST-mediated multifunctional RNA modulation. HEK293T cells were co-transfected with ddCas13b and MCP/PUFc-fused effectors. Desired manipulation consequences were achieved by addition of gRNAs tagged by MS2 or PBSc. All CREST components used in each condition were listed in the middle. (A) Simultaneous induction of alternative splicing by gRNA-1×MS2 and A-to-G base editing by gRNA-3×PBSc-Loop. Top: Inclusion of exon 7 in *SMN2* transcript measured by RT-qPCR. Middle: The design of transfection groups. Bottom: A-to-G editing efficiency quantified by sanger sequencing. (n=3) (B) Simultaneous induction of alternative splicing by gRNA-1×MS2 and C-to-U base editing by gRNA-10xPBSc-Loop. Top: Inclusion of exon 7 in *SMN2* transcript measured by RT-qPCR. Middle: The design of transfection groups. Bottom: C-to-U editing efficiency quantified by sanger sequencing. (n=3) (C) Simultaneous A-to-G base editing by gRNA-1xMS2 and C-to-U based editing by gRNA-10×PBSc-Loop. A-to-G (Top) and C-to-U (Bottom) editing efficiency quantified by sanger sequencing. (n=2). Data were displayed as mean ± S.E.M. *P<0.05, ** P < 0.01, ***P<0.001, ****P<0.0001, ns, not significant, by two-sided t-test.

## Discussion

RNA bridges the genetic information flow from DNA to protein, but also functions independent of translation as non-coding RNAs(39, 40). Given their importance, RNA-centric methodologies have been explored for decades and advanced significantly by the latest discovery of the CRISPR-Cas13 system. Compared to genomic editing, transcriptome engineering offers several advantages. Firstly, it does not introduce permanent, heritable changes. Secondly, unlike CRISPR-Cas9 systems that require a protospacer adjacent motif (PAM) for DNA targeting, the protospacer flanking site (PFS) sequence for Cas13 is more flexible(18). Last but not least, some diseases, such as m6A methylation-related cancers, are caused by altered RNA modification and processing that cannot be directly corrected by genome editing(41). However, the current RNA-targeting molecular toolkit does not meet the demand of simultaneously engineering multiple transcripts at different regulatory layers.

In this study, we filled this gap by coupling CRISPR-Cas13 with RNA scaffold systems. Our novel platform, called CREST, is compatible with at least three distinct functions, alternative splicing modulation, A-to-G and C-to-U base editing. By rational design of the linker between PBSc, we significantly improved the efficiency of CREST-PUFc-RBFOX1-mediated exon 7 inclusion in a SMN2 minigene transcript, revealing the additive effect of alternative splicing regulator RBFOX1. However, no significant difference was observed between 3 and 10 copies of PBSc with loop on the CREST-PUFc-ADAR2-DD-mediated base editing efficiency. This difference indicates that our CREST platform can be used to study the regulatory kinetics of RNA binding proteins and enzymes. Though tandem MS2 scaffolds have been optimized for signal amplification in genome imaging(42), it needs to be further investigated for RNA engineering. By protein alignment between PguCas13b and PspCas13b and mutational analysis, we identified that the K367 of PspCas13b is likely required for gRNA processing abrogation of which by K367A mutation and better yet the additional D369A and K370A mutations allows the appended gRNA scaffolds to be retained on gRNA for CREST functionality.

On top of that, we demonstrated the orthogonality of different RNA scaffolds in our CREST platform and concurrently multifunctional RNA modulation for the first time. Enabled by CREST, we simultaneously modulated alternative splicing and base editing by providing function-matching gRNAs that can specifically recruit corresponding effectors at the respective targets. CREST ddCas13b and RBD-effectors can be “pre-installed” into the cells to which the addition of gRNAs alone can induce combinatorial editing and modulations of multiple different transcripts.

An additional application of our CREST system is its utility in potentially reducing dCas13-indpendent off-target effects encountered when full-length effectors are utilized (17). Restoring enzymatic activity of split ADAR at desired locus was shown to be an effective strategy to reduce off-target effect on both RNA and DNA base editing (37, 43). We demonstrated the reconstitution of ADAR2-DD from two split halves using CREST architecture(37). Strikingly, compared with the full length PUFc-ADAR2-DD, our CREST system can reconstitute near 80% of catalytical activity of split ADAR2-DD and decrease about 96% of off-target events. We see potential of this CREST-mediated reconstitution strategy for other effectors and enzymes. For example, by splitting and reconstituting engineered ascorbate peroxidase enzyme (APEX2) in proximity RNA labelling approaches to characterize the RNA-bound proteome utilizing our CREST system may improve the reproducibility and reduce the background of random labeling(44, 45). We foresee that CREST can act like a socket with flexible functions and broad applications in RNA biology field.

In sum, we developed a new method called CREST to overcome the challenges in CRISPR-Cas13 mediated RNA engineering. By tagging gRNAs with orthogonal RNA scaffolds, CREST fills the gap of simultaneously modulating multiple transcripts for different functions, such as alternative splicing and base editing. This architecture allows the introduction of additional functions into the RNA modulation toolbox via orthogonal scaffolds and effectors. In addition, CREST enables us to reconstitute the enzymatic activity of split ADAR2 with high efficiency and specificity. Overall, CREST system may benefit a wide range of applications in RNA-centric research and therapeutic development.

## METHODS

### Plasmid construction

All coding sequences used in this study were cloned into pmax expression vector (Lonza) by TEDA as previously reported (46). IDT gBlocks encoding fragments of dPspCas13b as in pC0039-CMV-dPspCas13b-GS-ADAR2DD(E488Q) (Addgene# 103849) (25) were used as PCR templates for generating dPspCas13 and its variants. All mutations were designed in the primers and introduced to constructs by PCR and TEDA. An IDT gBlock encoding ADARDD(E488Q) sequence as in pC0039-CMV-dPspCas13b-GS-ADAR2DD(E488Q) (Addgene# 103849) (25) was used as PCR template for split ADAR2-DD and other ADAR2-DD fusion proteins. For split version, ADAR2-DD was split into two fragments between amino acid residues E466 and P467 with PCR and added into ddCas13 or PUFc by TEDA. pC0079-pCMV-dCas13b6-mapkNES-GS-dADAR2 (Addgene #130662, a gift from Feng Zhang) (19) was used for split RESCUE-S and other RESCUE-S fusion proteins. The split site in RESCUE-S remained the same as ADAR2-DD. pCI-SMN2 (Addgene #72287, a gift from Elliot Androphy) containing genomic region from exon 6 to 8 of *SMN2* gene placed downstream of a CMV promoter served as splicing minigene. For A-to-G base editing mScarlet reporter, a premature stop codon (UAG) was introduced into the coding region of mScarlet by PCR to enable the easy readout with flow cytometry. For C-to-U base editing reporter, we ordered the oligos with the previous reported targeting region ‘CTGATCTGCCTGTCCCACATCAATCGA’ and inserted it into the downstream of mScarlet(19) by annealing and ligation. For human diseases related genes, we selected multiple genes for both A-to-G and C-to-U editing, designed the gBlock (IDT) by copying the surrounding 190nt near the mutation site in each gene, and inserted it into the downstream of CMV promoter by PCR and TEDA ligation.

gRNA expression cloning plasmids were cloned into a pCR8 vector with U6 promoter and a ccdbCam selection cassette flanked by two BbsI restriction enzyme cut sites as described previously(20). Oligos of spacer sequence in gRNAs were designed manually and ordered from IDT (Supplementary Table S2). Oligos were annealed and ligated into BbsI-digested gRNA backbone. To append RNA scaffolds to 3’ of gRNA, gRNA backbone was digested by BsaI and Bglll and then ligated with annealed oligos containing RNA scaffold (Supplementary Table S2).

### Cell culture and transfection

HEK293T (ATCC) cells were grown in Dulbecco’s modified Eagle’s medium (DMEM) (Sigma) with 10% fetal bovine serum (FBS)(ThermoFisher), 4% Glutamax (ThermoFisher), 1% Sodium Pyruvate (ThermoFisher) and penicillin-streptomycin (ThermoFisher). Incubator conditions were 37°C and 5% CO2. For transfection, HEK293T cells were seeded in 12-well, 24-well or 96-well plates at the density of 1.5-2 million cells per plate one day prior to transfection. All transfection experiments were done with Lipo3000 (ThermoFisher) according to manufacturer’s instructions. The total amount and ratio of plasmids in each experiment can be found in Supplementary Table S1. Cells were harvested 48 h after transfection for flow cytometry or RNA extraction.

### RT-qPCR and Sanger sequencing

HEK293T cells were harvested for RNA extraction using RNeasy Plus Mini Kit (Qiagene) or NucleoSpin RNA Plus Kit (Takara). RNA concentration was measured by nanodrop and 500ng to 1000ng total RNAs were used for reverse transcription in 10 ul reaction volume by High-Capacity RNA-to-cDNA kit (ThermoFisher). Equal amount of RNAs were used in the same experiment.

For qPCR, 4 ul of diluted cDNA (1:40 dilution rate) were used for each reaction with 5 ul SsoAdvanced Universal SYBR® Green Supermix (BIO-RAD) and 1 ul of primer mix (5uM). qPCR reaction was run in ViiA 7 Real-Time PCR System (ThermoFisher) with standard program. The inclusion/exclusion ratio was calculated by 2~(Ct_-exclusion_ – Ct_-inclusion_) and then normalized to the condition of control gRNA in the same experiment (displayed as first column in each figure) to determine the fold change of inclusion/exclusion ratio. All primer sequences are listed in Supplementary Table S2.

For sanger sequencing, 2 ul of cDNA without dilution was used for each PCR reaction by Phusion® HighFidelity DNA Polymerase (NEB) for 35 cycles. PCR products were loaded on a 1% agarose gel, run at 140 Voltage for 50 mins, and purified by QIAquick PCR Purification Kit (Qiagene). Sanger sequencing was done in Quintarabio at UConn Health. All primer sequences were listed in Supplementary Table S2.

### Quantification of base editing efficiency

Data (.ab1 format) from sanger sequencing were uploaded to the online tool EditR for quantification (https://moriaritylab.shinyapps.io/editr_v10/ (38). 100 nucleotides around the target base with clean sequencing peaks were used to normalize signal. 30 nucleotides spanning the target base was used as query sequence as indicated in Figure 3C and EditR scored the percent area of the signal for each base (ACGT) at each position along the query sequence. The editing efficiency was calculated as AtoG=G_-score_/(A_-score_+G_-score_)×100% and CtoU=T_-score_/(C_-score_+T_-score_)×100%.

### Flow cytometry

Cells were seeded and transfected in a 96-well plate. For cell collection, media were discarded and 40 ul of trypsin (0.25%) were added into each well for incubation (8 mins at 37 °C). Then 80 ul of fresh media were directly added on the top of trypsin and mixed by pipetting up and down. All cells were transferred to a V bottom 96-well plate to run flow cytometry analysis on LSRFortessa X-20 (BD Bioscience). mScarlet / Clover ratio was calculated as Q2/(Q2+Q4) based on the percentage of each gate (Supplementary Figure S3).

### RNA isolation and next-generation sequencing

HEK293T cells were transfected with given plasmids and then harvested for RNA extraction by NucleoSpin RNA Plus Kit (Takara) 48 hour after transfection. All RNAs were sent to GENEWIZ for next generation sequencing and cDNA libraries were generated by polyA selection and yielded 24.3 million reads on average for each sample. All reads were mapped to hg38 by nextflow-core pipeline to get the .bam file as input for REDItool to quantify A-to-G base editing efficiency. Significant editing events were identified by get_DE_events.py in REDItool (47).

## ACKNOWLEDGEMENT

We thank the Flow Cytometry Service Core Facility at the Jackson Laboratory for providing technical support. We thank all the lab members in Cheng Lab for experimental support and instructive discussions.

