## Supplementary Information for "Simultaneous multifunctional transcriptome engineering by CRISPR RNA scaffold"

**Supplementary Figures S1-S9**

**Supplementary Protein Sequences**

**Supplementary Table S1-S2**

### Supplementary Figures

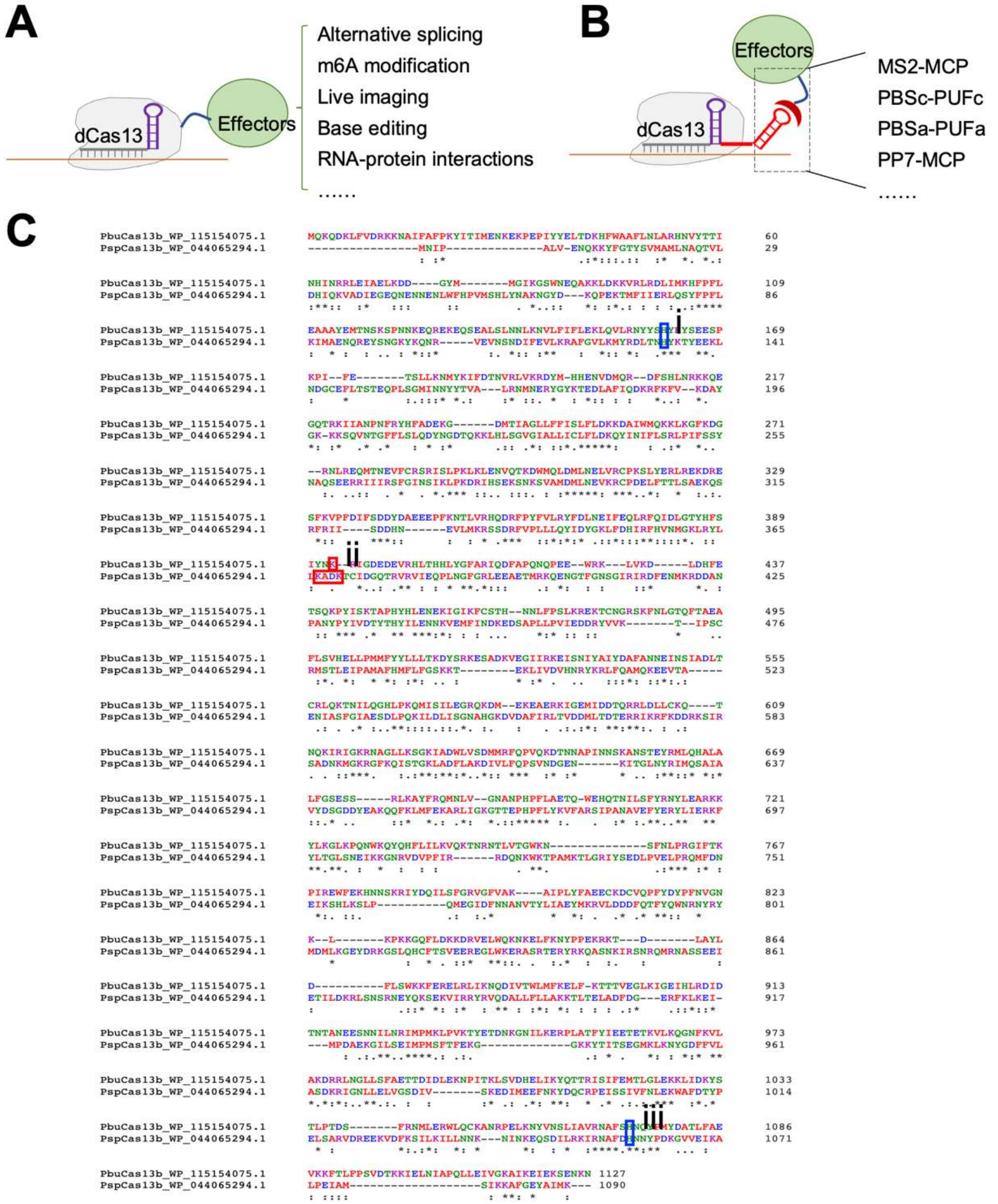

**Supplemental figures S1. Combinatorial RNA Editing via Scaffold Tagged gRNA** (A) Conventional CRISPR/Cas13 mediated RNA editing. gRNA consists of two parts, the spacer (gray) to match with targeting transcripts and the direct repeats (DR, purple) to bind with Cas13 protein. Different effector can be fused with dCas13 to execute different functions as indicated. (B) In CREST system, one or multiple copies of scaffold RNA motif (red) are added to the 3' end of gRNA and can be recognized and bound by specific RNA binding domains (RBDs). Effectors are fused with RBDs for RNA manipulation. Some scaffold RNA motifs and their cognate RBDs are listed at the right. (C) Alignment of PbuCas13b and PspCas13b. The amino acids in blue rectangle are mutated to alanine to generate the dCas13 and the amino acids in red rectangle are mutated to alanine to disable the crRNA processing activity of Cas13 in the ddCas13b.

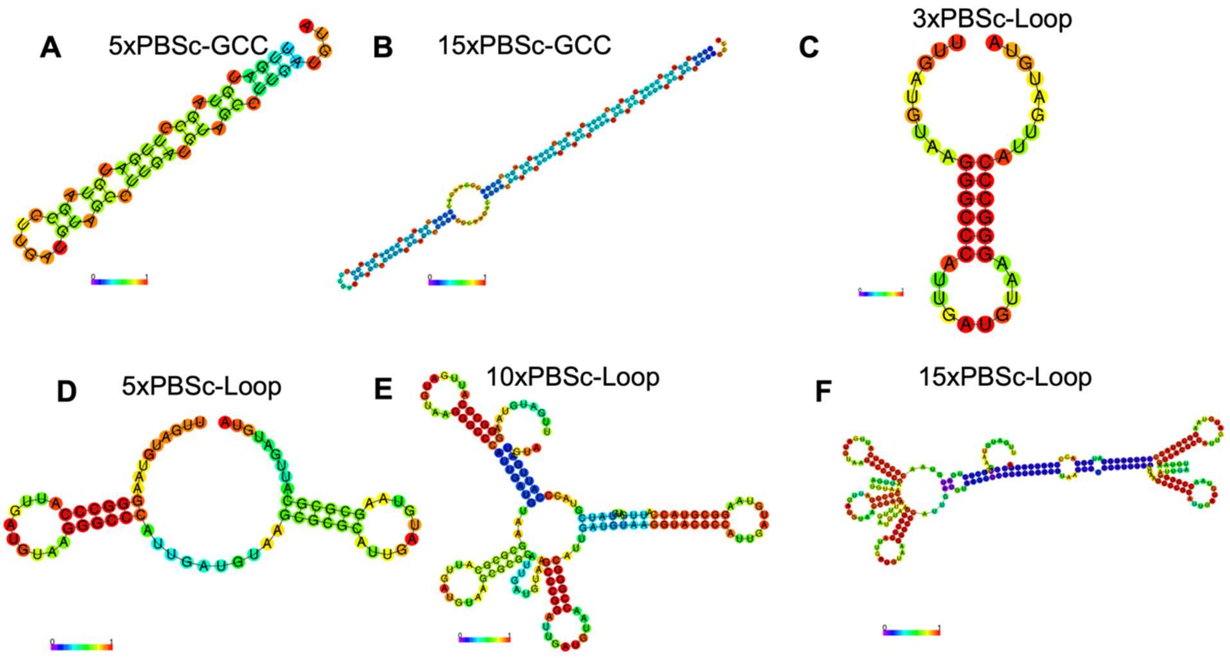

**Supplemental figures S2 Predicted secondary structures of PBSc scaffolds by online tool RNAfold.** (A-B) 5 and 15 copies of PBSc with GCC linker. The sequence of PBSc is UUGAUGUA. (C-F) Different copies of PBSc stabilized by high GC content stem-loops. 3/5/10/15 copies from C-F separately.

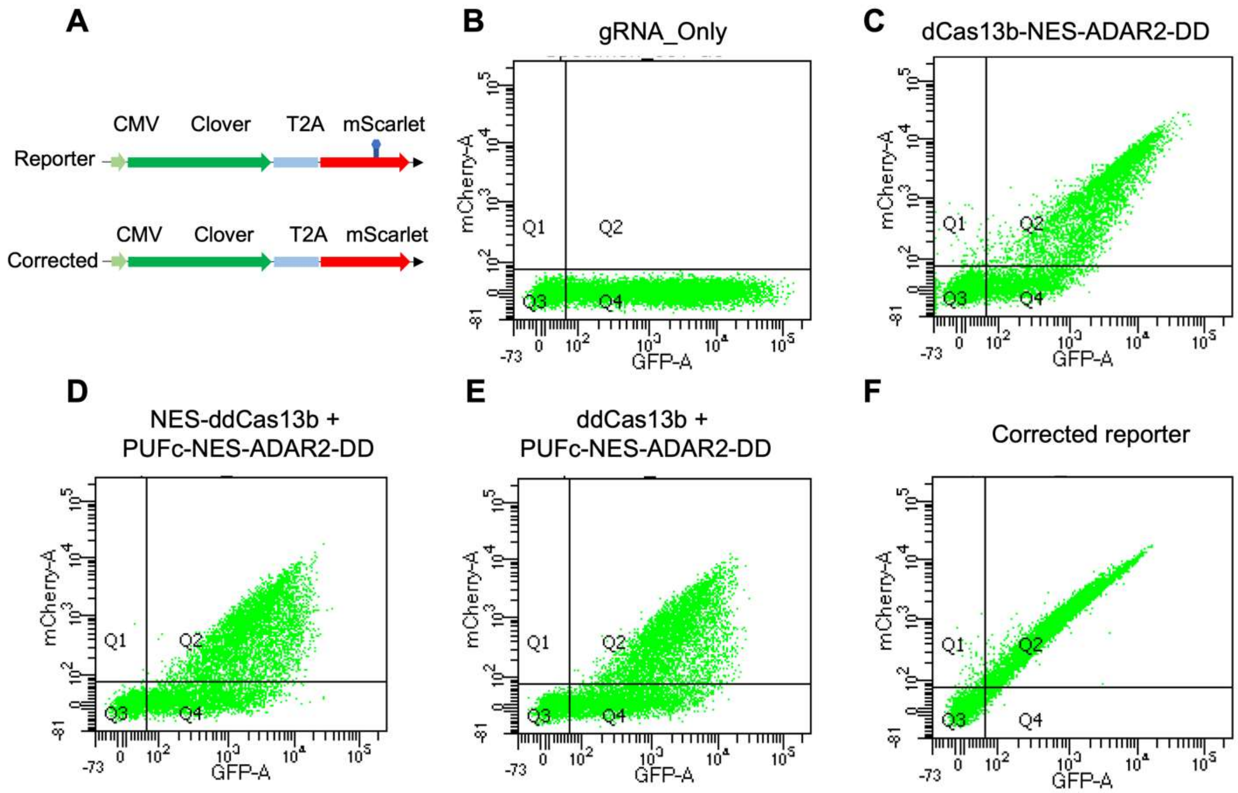

**Supplementary figure S3 Representative flow cytometry plots of A-to-G editing by CREST**. (A) Diagram of reporter construct for A-to-G editing with a premature stop codon in mScarlet and the corrected reporter without mutation. (B-E) HEK293T cells were co-transfected with reporter minigene harboring premature stop codon in the coding region of mScarlet and CREST components for correcting this mutation as indicated on the top. (F) HEK293T cells were transfected with reporter minigene without premature stop codon in mScarlet as a positive control.

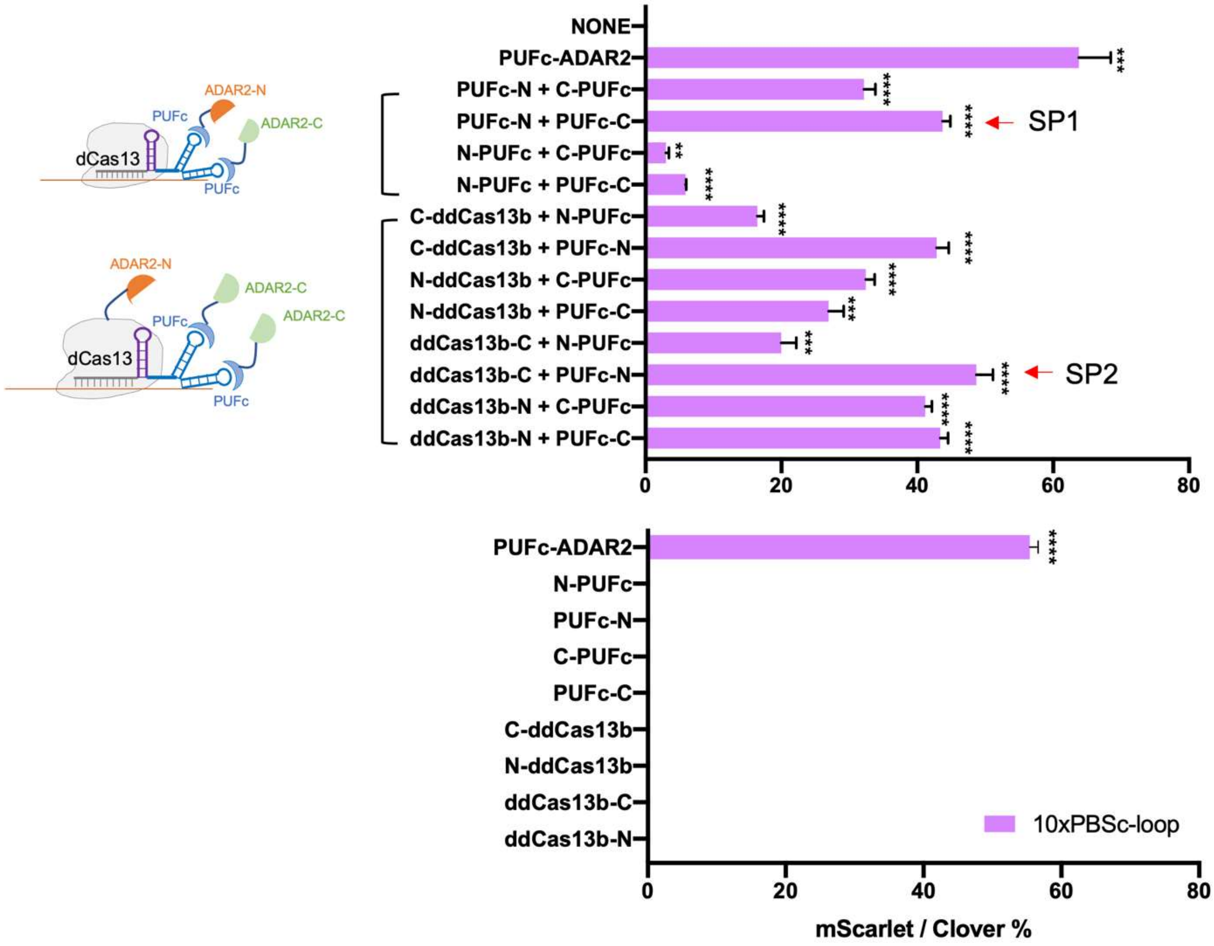

**Supplementary figures S4. Half of split ADAR2-DD is insufficient for A-to-G editing**. Top: HEK293T cells were transfected with different combinations of split ADAR2-DD and gRNA tagged with 10xPBSc loop. Bottom: HEK293T cells were transfected with single part of split ADAR2-DD and showed no activity for A-to-G editing. Data were displayed as mean ± S.E.M, n = 3. *P<0.05, ** P < 0.01,***P<0.001, ****P<0.0001, ns, not significant, by t-test.

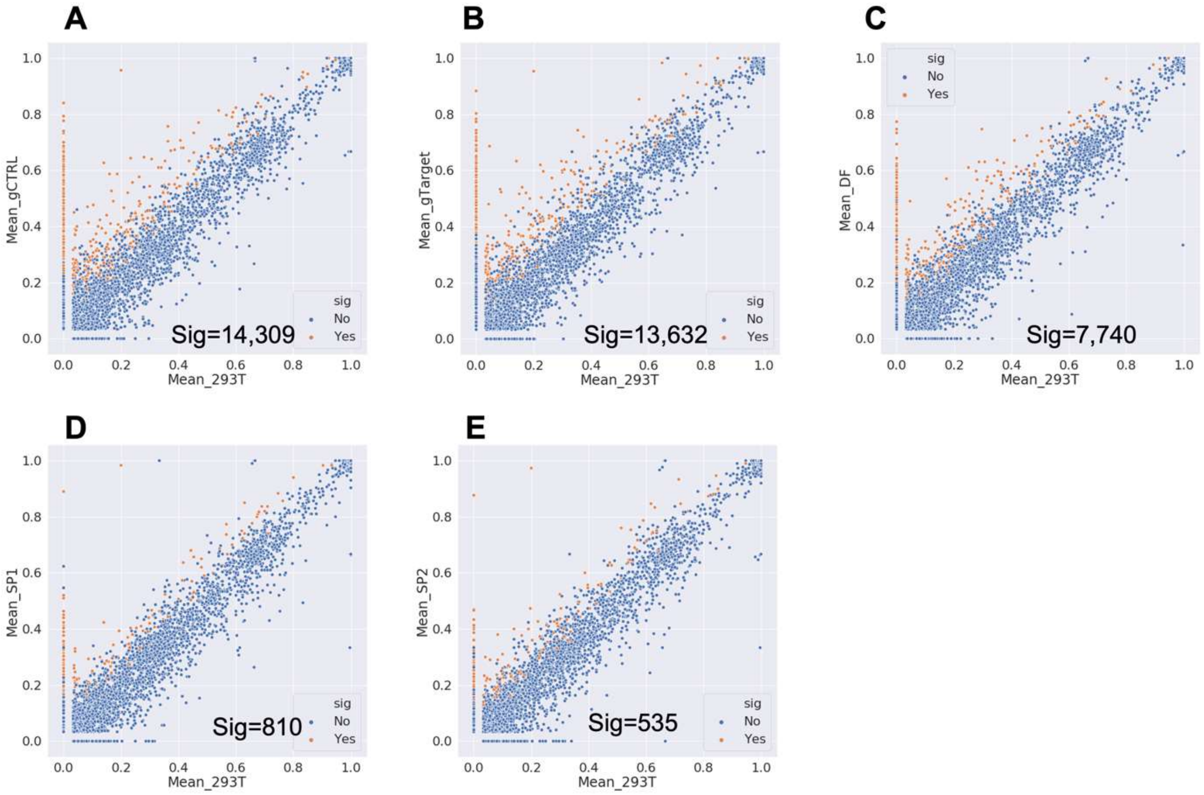

**Supplemental figures S5. Transcriptome-wide off-target analysis of A-to-G editing.** 2D scatter plot comparing the A-to-G editing yields observed with each construct (y-axis) to the yields observed with the control sample (x-axis, plain 293T). (A-B)gCtrl and gTarget stand for HEK293T cells co-transfected with ddCas13, PUFc-ADAR2-DD and indicated gRNA respectively. (C) HEK293T cells transfected with direct fusion of dCas13-ADAR2-DD. (E-F) Reconstitution of split ADAR2-DD activity by SP1 and SP2. Dots highlighted in red stand for sites with significant changes in A-to-G editing yields when comparing treatment to plain 293T without transfection. The number of significant editing sites (Sig) is listed in each comparison at the bottom right.

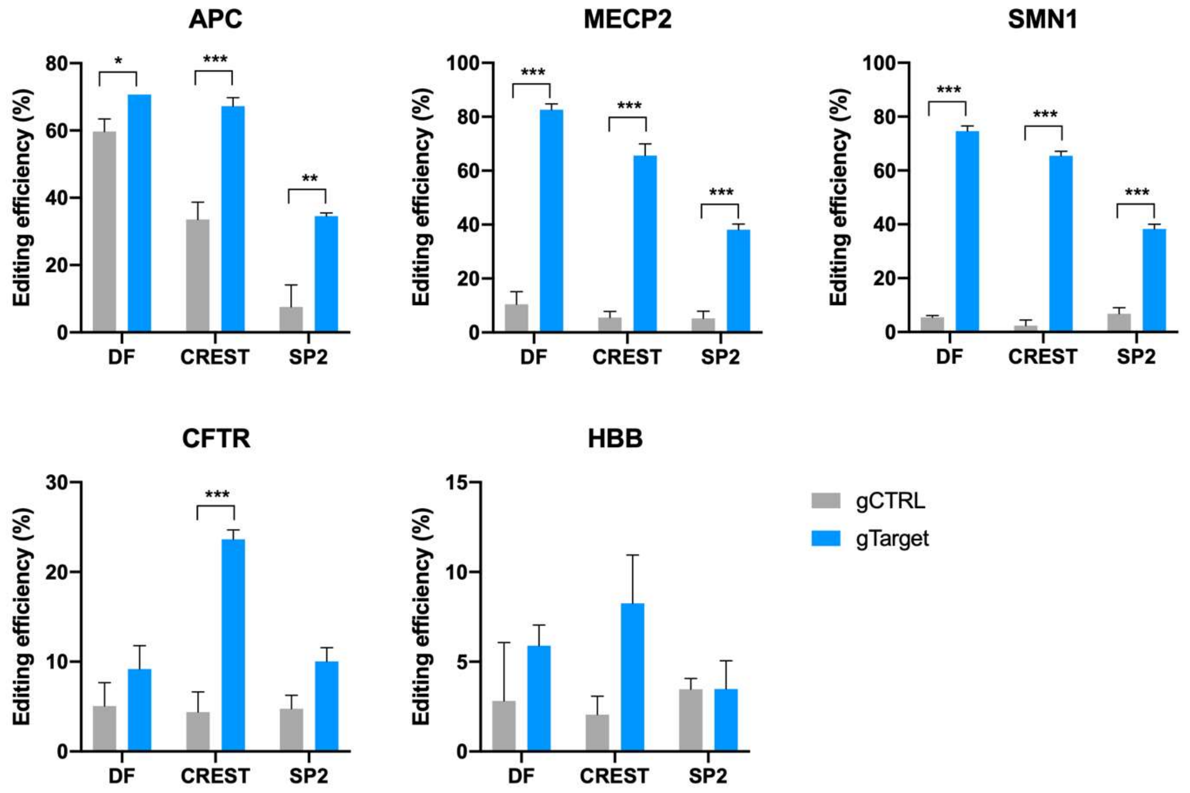

**Supplemental figures S6. A-to-G editing of disease-relevant mutations on reporters by CREST system.** Editing efficiency was measured by RT-PCR followed sanger sequencing (Y axis) and different treatments were annotated as X axis. DF: dCas13-ADAR2-DD. CREST: ddCas13+PUFc-ADAR2-DD. SP2: ddCas13b-ADAR2-DD-C + PUFc-ADAR2-DD-N. All gene names were indicated on top of each graph, gray stands for non-target control gRNA and blue stands for on-target gRNA. All gRNAs were tagged by 3xPBSc with mismatch distance of 22. Data were displayed as mean ± S.E.M, n = 3. *P<0.05, ** P < 0.01,***P<0.001, by t-test.

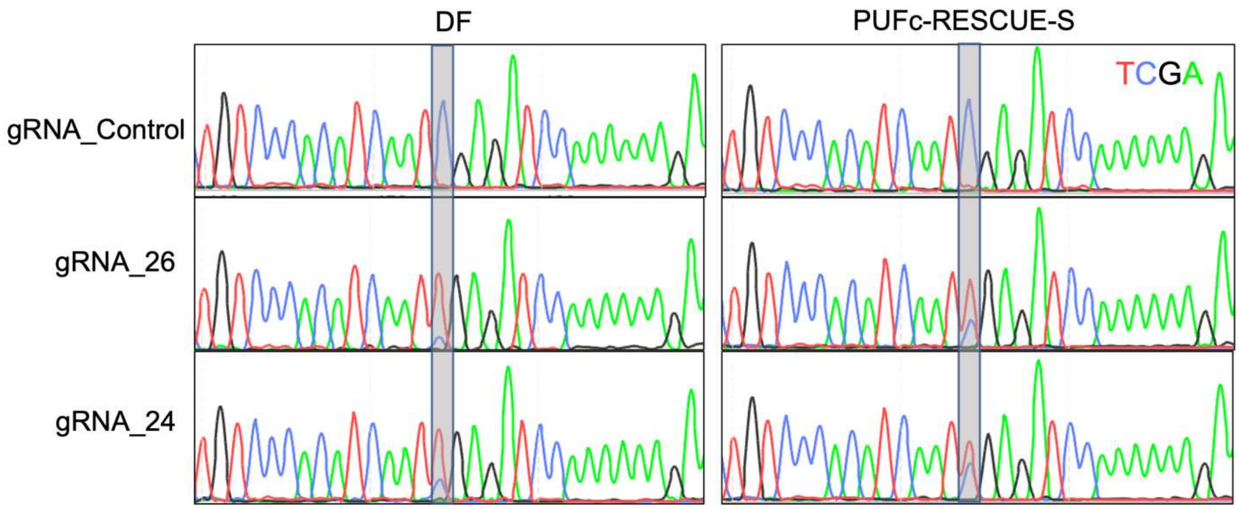

**Supplemental figures S7. Representative electropherograms showing Sanger sequencing results of C-to-U base editing.** Non-target control and on-target gRNA with 24nt and 26nt of mismatch distance were showed. DF stands for dCas13-RESCUE-S. Targeting site was highlighted in gray and colors for different bases were annotated on the top right.

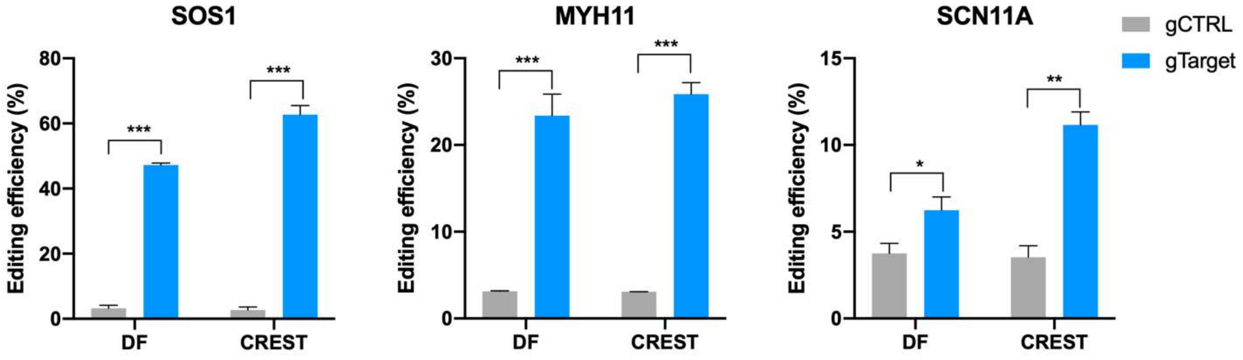

**Supplemental figures S8. C-to-U editing of disease-relevant mutations on reporters by CREST system.** Editing efficiency was measured by RT-PCR followed sanger sequencing (Y axis) and different treatments were annotated as X axis. DF: dCas13-ADAR2-DD. CREST: ddCas13+PUFc-RESCUE-S. All gene names were indicated on top of each graph, gray stands for non-target control gRNA and blue stands for on-target gRNA. All gRNAs were tagged by 3xPBSc with mismatch distance of 22. Data were displayed as mean ± S.E.M, n = 3. *P<0.05, ** P < 0.01,***P<0.001, by t-test.

```
ADAR2-DD      QLHLPQVLADAVSRLVLGKFGDLTDNFSSPHARRKVLAGVVMTTGTDVKDAKVISVSTGT
RESCUE-S      QLHLPQVLADAVSRLVIGKFGDLTDNFSSPHARRIGLAGVVMTTGTDVKDAKVICVSTGA
              ****************:*****************  ******************.****:

ADAR2-DD      KCINGEYMSDRGLALNDCHAEIISRRSLLRFLYTQLELYLNNKDDQKRSIFQKSERGGFR
RESCUE-S      KCINGEYLSDRGLALNDCHAEIVSRRSLLRFLYTQLELYLNNEDDQKRSIFQKSERGGFR
              *******:**************:*******************:*****************

ADAR2-DD      LKENVQFHLYISTSPCGDARIFSPHEPILEEPADRHPNRKARGQLRTKIESGQGTIPVRS
RESCUE-S      LKENIQFHLYISTSPCGDARIFSPHEAILEEPADRHPNRKARGQLRTKIEAGQGTIPVRN
              ****:********************* ***********************:********.

ADAR2-DD      NASIQTWDGVLQGERLLTMSCSDKIARWNVVGIQGSLLSIFVEPIYFSSIILGSLYHGDH
RESCUE-S      NASIQTWDGVLQGERLLTMSCSDKIARWNVVGIQGSLLSIFVEPIYFSSIILGSLYHGDH
              ************************************************************

ADAR2-DD      LSRAMYQRISNIEDLPPLYTLNKPLLSGISNAEARQPGKAPNFSVNWTVGDSAIEVINAT
RESCUE-S      LSRAMYQRISNIEDLPPLYTLNKPLLTGISNAEARQPGKAPIFSVNWTVGDSAIEVINAT
              **************************:************** ******************

ADAR2-DD      TGKDELGRASRLCKHALYCRWMRVHGKVPSHLLRSKITKPNVYHESKLAAKEYQAAKARL
RESCUE-S      TGKGELGRASRLCKHALYCRWMRVHGKVPSHLLRSKITKPNVYHETKLAAKEYQAAKARL
              ***.*****************************************:**************

ADAR2-DD      FTAFIKAGLGAWVEKPTEQDQFSLT 385
RESCUE-S      FTAFIKAGLGAWVEKPTEQDQFSLT 385
              *************************
```

**Supplemental figures S9.** Alignment of ADAR2-DD and RESCUE-S protein sequences. The conserved motif for splitting was highlighted in red and split site was indicated by red arrow between E and P.

**Supplementary Protein Sequences**

3xFLAG-ddpspCas13b:

MDYKDHDGDYKDHDIDYKDDDDKIDGGGGSDRRAMNIPALVENQKKYFGTYSVMAMLNAQTVLDHIQK
VADIEGEQNENNENLWFHPVMSHLYNAKNGYDKQPEKTMFIIERLQSYFPFLKIMAENQREYSNGKYKQ
NRVEVNSNDIFEVLKRAFGVLKMYRDLTNAYKTYEEKLNDGCEFLTSTEQPLSGMINNYYTVALRNMNER
YGYKTEDLAFIQDKRFKFVKDAYGKKKSQVNTGFFLSLQDYNGDTQKKLHLSGVGIALLICLFLDKQYINIF
LSRLPIFSSYNAQSEERRIIIRSFGINSIKLPKDRIHSEKSNKSVAMDMLNEVKRCPDELFTTLSAEKQSRFR
IISDDHNEVLMKRSSDRFVPLLLQYIDYGKLFDHIRFHVNMGKLRYLLAAAATCIDGQTRVRVIEQPLNGFG
RLEEAETMRKQENGTFGNSGIRIRDFENMKRDDANPANYPYIVDTYTHYILENNKVEMFINDKEDSAPLLP
VIEDDRYVVKTIPSCRMSTLEIPAMAFHMFLFGSKKTEKLIVDVHNRYKRLFQAMQKEEVTAENIASFGIAE
SDLPQKILDLISGNAHGKDVDAFIRLTVDDMLTDTERRIKRFKDDRKSIRSADNKMGKRGFKQISTGKLAD
FLAKDIVLFQPSVNDGENKITGLNYRIMQSAIAVYDSGDDYEAKQQFKLMFEKARLIGKGTTEPHPFLYKV
FARSIPANAVEFYERYLIERKFYLTGLSNEIKKGNRVDVPFIRRDQNKWKTPAMKTLGRIYSEDLPVELPR
QMFDNEIKSHLKSLPQMEGIDFNNANVTYLIAEYMKRVLDDDFQTFYQWNRNYRYMDMLKGEYDRKGSL
QHCFTSVEEREGLWKERASRTERYRKQASNKIRSNRQMRNASSEEIETILDKRLSNSRNEYQKSEKVIRR
YRVQDALLFLLAKKTLTELADFDGERFKLKEIMPDAEKGILSEIMPMSFTFEKGGKKYTITSEGMKLKNYGD
FFVLASDKRIGNLLELVGSDIVSKEDIMEEFNKYDQCRPEISSIVFNLEKWAFDTYPELSARVDREEKVDFK
SILKILLNNKNINKEQSDILRKIRNAFDANNYPDKGVVEIKALPEIAMSIKKAFGEYAIMKGGRGGGGSGGG
GSGGGGSGPA

hcoRBFOX1N-MCP-RBFOX1C

MNCEREQLRGNQEAAAAPDTMAQPYASAQFAPPQNGIPAEYTAPHPHPAPEYTGQTTVPEHTLNLYPP
AQTHSEQSPADTSAQTVSGTATQTDDAAPTDGQPQTQPSENTENKSQPKGGGGSGRAMASNFTQFVL
VDNGGTGDVTVAPSNFANGVAEWISSNSRSQAYKVTCSVRQSSAQKRKYTIKVEVPKVATQTVGGVELP
VAAWRSYLNMELTIPIFATNSDCELIVKAMQGLLKDGNPIPSAIAANSGIYSAGGRGGGGSGGGGSGGGG
SGPANATARVMTNKKTVNPYTNGWKLNPVVGAVYSPEFYAGTVLLCQANQEGSSMYSAPSSLVYTSAM
PGFPYPAATAAAAYRGAHLRGRGRTVYNTFRAAAPPPPIPAYGGVVYQDGFYGADIYGGYAAYRYAQPT

PATAAAYSDSYGRVYAADPYHHALAPAPTYGVGAMNAFAPLTDAKTRSHADDVGLVLSSLQASIYRGGY
NRFAPY

hcoRBFOX1N-PUFc-2xNLS-RBFOX1C

MNCEREQLRGNQEAAAAPDTMAQPYASAQFAPPQNGIPAEYTAPHPHPAPEYTGQTTVPEHTLNLYPP
AQTHSEQSPADTSAQTVSGTATQTDDAAPTDGQPQTQPSENTENKSQPKGGGGSGRAGILPPKKKRKV
SRGRSRLLEDFRNNRYPNLQLREIAGHIMEFSQDQHGSRFIQLKLERATPAERQLVFNEILQAAYQLMVD
VFGNYVIQKFFEFGSLEQKLALAERIRGHVLSLALQMYGSRVIEKALEFIPSDQQNEMVRELDGHVLKCVK
DQNGNHVVQKCIECVQPQSLQFIIDAFKGQVFALSTHPYGCRVIQRILEHCLPDQTLPILEELHQHTEQLV
QDQYGSYVIEHVLEHGRPEDKSKIVAEIRGNVLVLSQHKFANNVVQKCVTHASRTERAVLIDEVCTMNDG
PHSALYTMMKDQYANYVVQKMIDVAEPGQRKIVMHKIRPHIATLRKYTYGKHILAKLEKYYMKNGVDLGD
PKKKRKVDPKKKRKVGGRGGGGSGGGGSGGGGSGPANATARVMTNKKTVNPYTNGWKLNPVVGAVY
SPEFYAGTVLLCQANQEGSSMYSAPSSLVYTSAMPGFPYPAATAAAAYRGAHLRGRGRTVYNTFRAAA
PPPPIPAYGGVVYQDGFYGADIYGGYAAYRYAQPTPATAAAYSDSYGRVYAADPYHHALAPAPTYGVGA
MNAFAPLTDAKTRSHADDVGLVLSSLQASIYRGGYNRFAPY

MCP-NES-ADAR2DD(E448Q)

MASNFTQFVLVDNGGTGDVTVAPSNFANGVAEWISSNSRSQAYKVTCSVRQSSAQKRKYTIKVEVPKVA
TQTVGGVELPVAAWRSYLNMELTIPIFATNSDCELIVKAMQGLLKDGNPIPSAIAANSGIYSAGGRGGGGS
GGGGSGGGGSGPALQLPPLERLTLGSGGGGSQLHLPQVLADAVSRLVLGKFGDLTDNFSSPHARRKVL
AGVVMTTGTDVKDAKVISVSTGTKCINGEYMSDRGLALNDCHAEIISRRSLLRFLYTQLELYLNNKDDQKR
SIFQKSERGGFRLKENVQFHLYISTSPCGDARIFSPHEPILEEPADRHPNRKARGQLRTKIESGQGTIPVR
SNASIQTWDGVLQGERLLTMSCSDKIARWNVVGIQGSLLSIFVEPIYFSSIILGSLYHGDHLSRAMYQRISNI
EDLPPLYTLNKPLLSGISNAEARQPGKAPNFSVNWTVGDSAIEVINATTGKDELGRASRLCKHALYCRWM
RVHGKVPSHLLRSKITKPNVYHESKLAAKEYQAAKARLFTAFIKAGLGAWVEKPTEQDQFSLTNV

PUFc-NES-ADAR2DD(E488Q)

MNVGGGGSGGGGSGGGGSGRASRGRSRLLEDFRNNRYPNLQLREIAGHIMEFSQDQHGSRFIQLKLE
RATPAERQLVFNEILQAAYQLMVDVFGNYVIQKFFEFGSLEQKLALAERIRGHVLSLALQMYGSRVIEKAL
EFIPSDQQNEMVRELDGHVLKCVKDQNGNHVVQKCIECVQPQSLQFIIDAFKGQVFALSTHPYGCRVIQR
ILEHCLPDQTLPILEELHQHTEQLVQDQYGSYVIEHVLEHGRPEDKSKIVAEIRGNVLVLSQHKFANNVVQ
KCVTHASRTERAVLIDEVCTMNDGPHSALYTMMKDQYANYVVQKMIDVAEPGQRKIVMHKIRPHIATLRK
YTYGKHILAKLEKYYMKNGVDLGGGRGGGGSGGGGSGGGGSGPALQLPPLERLTLGSGGGGSQLHLP
QVLADAVSRLVLGKFGDLTDNFSSPHARRKVLAGVVMTTGTDVKDAKVISVSTGTKCINGEYMSDRGLAL
NDCHAEIISRRSLLRFLYTQLELYLNNKDDQKRSIFQKSERGGFRLKENVQFHLYISTSPCGDARIFSPHEP
ILEEPADRHPNRKARGQLRTKIESGQGTIPVRSNASIQTWDGVLQGERLLTMSCSDKIARWNVVGIQGSL
LSIFVEPIYFSSIILGSLYHGDHLSRAMYQRISNIEDLPPLYTLNKPLLSGISNAEARQPGKAPNFSVNWTVG
DSAIEVINATTGKDELGRASRLCKHALYCRWMRVHGKVPSHLLRSKITKPNVYHESKLAAKEYQAAKARL
FTAFIKAGLGAWVEKPTEQDQFSLTNV

PUFc-NES-RESCUE-S

MNVGGGGSGGGGSGGGGSGRASRGRSRLLEDFRNNRYPNLQLREIAGHIMEFSQDQHGSRFIQLKLE
RATPAERQLVFNEILQAAYQLMVDVFGNYVIQKFFEFGSLEQKLALAERIRGHVLSLALQMYGSRVIEKAL
EFIPSDQQNEMVRELDGHVLKCVKDQNGNHVVQKCIECVQPQSLQFIIDAFKGQVFALSTHPYGCRVIQR
ILEHCLPDQTLPILEELHQHTEQLVQDQYGSYVIEHVLEHGRPEDKSKIVAEIRGNVLVLSQHKFANNVVQ
KCVTHASRTERAVLIDEVCTMNDGPHSALYTMMKDQYANYVVQKMIDVAEPGQRKIVMHKIRPHIATLRK
YTYGKHILAKLEKYYMKNGVDLGGGRGGGGSGGGGSGGGGSGPALQKKLEELELGSQLHLPQVLADAV
SRLVIGKFGDLTDNFSSPHARRIGLAGVVMTTGTDVKDAKVICVSTGAKCINGEYLSDRGLALNDCHAEIV
SRRSLLRFLYTQLELYLNNEDDQKRSIFQKSERGGFRLKENIQFHLYISTSPCGDARIFSPHEAILEEPADR
HPNRKARGQLRTKIEAGQGTIPVRNNASIQTWDGVLQGERLLTMSCSDKIARWNVVGIQGSLLSIFVEPIY
FSSIILGSLYHGDHLSRAMYQRISNIEDLPPLYTLNKPLLTGISNAEARQPGKAPIFSVNWTVGDSAIEVINA
TTGKGELGRASRLCKHALYCRWMRVHGKVPSHLLRSKITKPNVYHETKLAAKEYQAAKARLFTAFIKAGL
GAWVEKPTEQDQFSLTVDGSGSGSLPPLERLTL

NES-PUFc-ADAR2DD(E488Q)-splitN

MLPPLERLTLGGGGSGRASRGRSRLLEDFRNNRYPNLQLREIAGHIMEFSQDQHGSRFIQLKLERATPAE
RQLVFNEILQAAYQLMVDVFGNYVIQKFFEFGSLEQKLALAERIRGHVLSLALQMYGSRVIEKALEFIPSDQ
QNEMVRELDGHVLKCVKDQNGNHVVQKCIECVQPQSLQFIIDAFKGQVFALSTHPYGCRVIQRILEHCLP
DQTLPILEELHQHTEQLVQDQYGSYVIEHVLEHGRPEDKSKIVAEIRGNVLVLSQHKFANNVVQKCVTHAS
RTERAVLIDEVCTMNDGPHSALYTMMKDQYANYVVQKMIDVAEPGQRKIVMHKIRPHIATLRKYTYGKHIL
AKLEKYYMKNGVDLGGGRGGGGSGGGGSGGGGSGPAQLHLPQVLADAVSRLVLGKFGDLTDNFSSPH
ARRKVLAGVVMTTGTDVKDAKVISVSTGTKCINGEYMSDRGLALNDCHAEIISRRSLLRFLYTQLELYLNN
KDDQKRSIFQKSERGGFRLKENVQFHLYISTSPCGDARIFSPHEPILEE

NES-PUFc-ADAR2DD(E488Q)-splitC

MLPPLERLTLGGGGSGRASRGRSRLLEDFRNNRYPNLQLREIAGHIMEFSQDQHGSRFIQLKLERATPAE
RQLVFNEILQAAYQLMVDVFGNYVIQKFFEFGSLEQKLALAERIRGHVLSLALQMYGSRVIEKALEFIPSDQ
QNEMVRELDGHVLKCVKDQNGNHVVQKCIECVQPQSLQFIIDAFKGQVFALSTHPYGCRVIQRILEHCLP
DQTLPILEELHQHTEQLVQDQYGSYVIEHVLEHGRPEDKSKIVAEIRGNVLVLSQHKFANNVVQKCVTHAS
RTERAVLIDEVCTMNDGPHSALYTMMKDQYANYVVQKMIDVAEPGQRKIVMHKIRPHIATLRKYTYGKHIL
AKLEKYYMKNGVDLGGGRGGGGSGGGGSGGGGSGPAPADRHPNRKARGQLRTKIESGQGTIPVRSNA
SIQTWDGVLQGERLLTMSCSDKIARWNVVGIQGSLLSIFVEPIYFSSIILGSLYHGDHLSRAMYQRISNIEDL
PPLYTLNKPLLSGISNAEARQPGKAPNFSVNWTVGDSAIEVINATTGKDELGRASRLCKHALYCRWMRVH
GKVPSHLLRSKITKPNVYHESKLAAKEYQAAKARLFTAFIKAGLGAWVEKPTEQDQFSLTNV

NES-ddPspCas13b-ARA2DD(E488Q)-SplitC

MLPPLERLTLGGGGSGRAMNIPALVENQKKYFGTYSVMAMLNAQTVLDHIQKVADIEGEQNENNENLWF
HPVMSHLYNAKNGYDKQPEKTMFIIERLQSYFPFLKIMAENQREYSNGKYKQNRVEVNSNDIFEVLKRAF
GVLKMYRDLTNAYKTYEEKLNDGCEFLTSTEQPLSGMINNYYTVALRNMNERYGYKTEDLAFIQDKRFKF
VKDAYGKKKSQVNTGFFLSLQDYNGDTQKKLHLSGVGIALLICLFLDKQYINIFLSRLPIFSSYNAQSEERRI
IIRSFGINSIKLPKDRIHSEKSNKSVAMDMLNEVKRCPDELFTTLSAEKQSRFRIISDDHNEVLMKRSSDRF
VPLLLQYIDYGKLFDHIRFHVNMGKLRYLLAAAATCIDGQTRVRVIEQPLNGFGRLEEAETMRKQENGTFG
NSGIRIRDFENMKRDDANPANYPYIVDTYTHYILENNKVEMFINDKEDSAPLLPVIEDDRYVVKTIPSCRMS

TLEIPAMAFHMFLFGSKKTEKLIVDVHNRYKRLFQAMQKEEVTAENIASFGIAESDLPQKILDLISGNAHGK

DVDAFIRLTVDDMLTDTERRIKRFKDDRKSIRSADNKMGKRGFKQISTGKLADFLAKDIVLFQPSVNDGEN

KITGLNYRIMQSAIAVYDSGDDYEAKQQFKLMFEKARLIGKGTTEPHPFLYKVFARSIPANAVEFYERYLIE

RKFYLTGLSNEIKKGNRVDVPFIRRDQNKWKTPAMKTLGRIYSEDLPVELPRQMFDNEIKSHLKSLPQME

GIDFNNANVTYLIAEYMKRVLDDDFQTFYQWNRNYRYMDMLKGEYDRKGSLQHCFTSVEEREGLWKER

ASRTERYRKQASNKIRSNRQMRNASSEEIETILDKRLSNSRNEYQKSEKVIRRYRVQDALLFLLAKKTLTE

LADFDGERFKLKEIMPDAEKGILSEIMPMSFTFEKGGKKYTITSEGMKLKNYGDFFVLASDKRIGNLLELVG

SDIVSKEDIMEEFNKYDQCRPEISSIVFNLEKWAFDTYPELSARVDREEKVDFKSILKILLNNKNINKEQSDI

LRKIRNAFDANNYPDKGVVEIKALPEIAMSIKKAFGEYAIMKGGRGGGGSGGGGSGGGGSGPAPADRHP

NRKARGQLRTKIESGQGTIPVRSNASIQTWDGVLQGERLLTMSCSDKIARWNVVGIQGSLLSIFVEPIYFS

SIILGSLYHGDHLSRAMYQRISNIEDLPPLYTLNKPLLSGISNAEARQPGKAPNFSVNWTVGDSAIEVINATT

GKDELGRASRLCKHALYCRWMRVHGKVPSHLLRSKITKPNVYHESKLAAKEYQAAKARLFTAFIKAGLGA

WVEKPTEQDQFSLTNV

NES-ddPspCas13b-RESCUE-S-splitN

MLPPLERLTLGGGGSGRAMNIPALVENQKKYFGTYSVMAMLNAQTVLDHIQKVADIEGEQNENNENLWF

HPVMSHLYNAKNGYDKQPEKTMFIIERLQSYFPFLKIMAENQREYSNGKYKQNRVEVNSNDIFEVLKRAF

GVLKMYRDLTNAYKTYEEKLNDGCEFLTSTEQPLSGMINNYYTVALRNMNERYGYKTEDLAFIQDKRFKF

VKDAYGKKKSQVNTGFFLSLQDYNGDTQKKLHLSGVGIALLICLFLDKQYINIFLSRLPIFSSYNAQSEERRI

IIRSFGINSIKLPKDRIHSEKSNKSVAMDMLNEVKRCPDELFTTLSAEKQSRFRIISDDHNEVLMKRSSDRF

VPLLLQYIDYGKLFDHIRFHVNMGKLRYLLAAAATCIDGQTRVRVIEQPLNGFGRLEEAETMRKQENGTFG

NSGIRIRDFENMKRDDANPANYPYIVDTYTHYILENNKVEMFINDKEDSAPLLPVIEDDRYVVKTIPSCRMS

TLEIPAMAFHMFLFGSKKTEKLIVDVHNRYKRLFQAMQKEEVTAENIASFGIAESDLPQKILDLISGNAHGK

DVDAFIRLTVDDMLTDTERRIKRFKDDRKSIRSADNKMGKRGFKQISTGKLADFLAKDIVLFQPSVNDGEN

KITGLNYRIMQSAIAVYDSGDDYEAKQQFKLMFEKARLIGKGTTEPHPFLYKVFARSIPANAVEFYERYLIE

RKFYLTGLSNEIKKGNRVDVPFIRRDQNKWKTPAMKTLGRIYSEDLPVELPRQMFDNEIKSHLKSLPQME

GIDFNNANVTYLIAEYMKRVLDDDFQTFYQWNRNYRYMDMLKGEYDRKGSLQHCFTSVEEREGLWKER

ASRTERYRKQASNKIRSNRQMRNASSEEIETILDKRLSNSRNEYQKSEKVIRRYRVQDALLFLLAKKTLTE

LADFDGERFKLKEIMPDAEKGILSEIMPMSFTFEKGGKKYTITSEGMKLKNYGDFFVLASDKRIGNLLELVG
SDIVSKEDIMEEFNKYDQCRPEISSIVFNLEKWAFDTYPELSARVDREEKVDFKSILKILLNNKNINKEQSDI
LRKIRNAFDANNYPDKGVVEIKALPEIAMSIKKAFGEYAIMKGGRGGGGSGGGGSGGGGSGPAQLHLPQ
VLADAVSRLVIGKFGDLTDNFSSPHARRIGLAGVVMTTGTDVKDAKVICVSTGAKCINGEYLSDRGLALND
CHAEIVSRRSLLRFLYTQLELYLNNEDDQKRSIFQKSERGGFRLKENIQFHLYISTSPCGDARIFSPHEAILE
E

NES-PUFc-RESCUE-S-splitC

MLPPLERLTLGGGGSGRASRGRSRLLEDFRNNRYPNLQLREIAGHIMEFSQDQHGSRFIQLKLERATPAE
RQLVFNEILQAAYQLMVDVFGNYVIQKFFEFGSLEQKLALAERIRGHVLSLALQMYGSRVIEKALEFIPSDQ
QNEMVRELDGHVLKCVKDQNGNHVVQKCIECVQPQSLQFIIDAFKGQVFALSTHPYGCRVIQRILEHCLP
DQTLPILEELHQHTEQLVQDQYGSYVIEHVLEHGRPEDKSKIVAEIRGNVLVLSQHKFANNVVQKCVTHAS
RTERAVLIDEVCTMNDGPHSALYTMMKDQYANYVVQKMIDVAEPGQRKIVMHKIRPHIATLRKYTYGKHIL
AKLEKYYMKNGVDLGGGRGGGGSGGGGSGGGGSGPAPADRHPNRKARGQLRTKIEAGQGTIPVRNNA
SIQTWDGVLQGERLLTMSCSDKIARWNVVGIQGSLLSIFVEPIYFSSIILGSLYHGDHLSRAMYQRISNIEDL
PPLYTLNKPLLTGISNAEARQPGKAPIFSVNWTVGDSAIEVINATTGKGELGRASRLCKHALYCRWMRVH
GKVPSHLLRSKITKPNVYHETKLAAKEYQAAKARLFTAFIKAGLGAWVEKPTEQDQFSLT

**Supplementary Table S1**

| | | | | |
|---|---|---|---|---|
| Figure 1. B&C | 12-wells plate | | | |
| constructs | ddCas13 | RBD-Effector | gRNA | PCI-SMN2 reporter |
| plasmid amount (ng) | 150 | 150 | 100* | 18 |
| *For alternative splicing, three gRNAs were used in all experiments with equally split amount. | | | | |
| Figure 1. D / G | 12-wells plate | | | |
| constructs | ddCas13 | RBD-Effector | gRNA | PCI-SMN2 reporter |
| plasmid amount (ng) | 300 | 100 | 200 | 18 |
| Figure 1E | 24-well plate | | | |
| constructs | ddCas13 | RBD-Effector | gRNA | PCI-SMN2 reporter |
| plasmid amount (ng) | 150 | 50 | 100 | 9 |
| Figure 1.F | 12-wells plate | | | |
| constructs | ddCas13 | RBD-Effector | gRNA | PCI-SMN2 reporter |
| plasmid amount (ng) | 300 | 300 | 300 | 18 |
| Figure 2.B/C | 96-well plate | | | |
| constructs | ddCas13 | RBD-Effector | gRNA | AtoG reporter |
| plasmid amount (ng) | 100ng | 33ng | 66ng | 7ng |
| Figure 2. D/E Figure 3. B/C/D Figure S6, S8 | 24 well plate | | | |
| constructs | ddCas13 | RBD-Effector | gRNA | AtoG reporter |
| plasmid amount (ng) | 300ng | 100ng* | 200ng | 10ng |
| *split into 50ng+50ng for split ADAR2/RESCUE-S | | | | |

| | | | | |
|---|---|---|---|---|
| Figure 4. A /B/C | 24-well plate | | | |
| constructs | ddCas13 | RBD-effector-<br>1+2 | gRNA-1 + 2 * | reporter 1+2 |
| plasmid amount (ng) | 250 | 75+75 | 75 +75 | 10+10 |
| *the '-' symbol in the figure stands for the addition of control gRNA | | | | |

**Supplementary Table S2**

| PCR Primers | |
|---|---|
| AtoG editing Forward | GGCTCCGAATTCACCGGTG |
| AtoG editing Reverse | CTTCTTCTGCATTACGGGGC |
| CtoU editing Forward | GTGGGAGCGCGTGATGAACT |
| CtoU editing Reverse | AAAGCTGGGTCGAATTCTTAATTAA |
| Sanger sequencing primer | |
| AtoG sequencing | GCCCGACAACCACTACCTGA |
| CtoU sequencing | TGGACATCACCTCCCACAACG |

| qRT-PCR primers | |
|---|---|
| SMN2-Forward | GCTCTTAAGGCTAGAGTACTTAATACGA |
| SMN2-Inclusion-Reverse | CTTCTTTTTGATTTTGTCTAAAACCCATATAATAG |
| SMN2-Exclusion-Reverse | CTCTATGCCAGCATTTCCATATAATAG |

| gRNA spacer sequence oligos | |
|---|---|
| Ctrl | ACTCAAAAGGAAGTGACAAGAA |
| SMN1-gRNA1 | GTAAGATTCACTTTCATAATGC |
| SMN1-gRNA2 | GTAGGGATGTAGATTAACCTTT |
| SMN1-gRNA3 | GCTGGTCTGCCTACTAGTGATA |
| mScarlet-A-to-G-2 | GACATGAACTGAGGGGACAGGATGTCCCA |

| | |
|---|---|
| mScarlet-A-to-G-4 | GATGAACTGAGGGGACAGGATGTCCCAGG |
| mScarlet-A-to-G-6 | GATGAACTGAGGGGACAGGATGTCCCAGGAG |
| mScarlet-A-to-G-8 | GAACTGAGGGGACAGGATGTCCCAGGAGAA |
| mScarlet-A-to-G-10 | GACTGAGGGGACAGGATGTCCCAGGAGAAGG |
| mScarlet-A-to-G-12 | GTGAGGGGACAGGATGTCCCAGGAGAAGGGC |
| mScarlet-A-to-G-14 | GAGGGGACAGGATGTCCCAGGAGAAGGGCAG |
| mScarlet-A-to-G-16 | GGGACAGGATGTCCCAGGAGAAGGGCAGGG |
| mScarlet-A-to-G-18 | GACAGGATGTCCCAGGAGAAGGGCAGGGGG |
| mScarlet-A-to-G-20 | GCAGGATGTCCCAGGAGAAGGGCAGGGGGCC |
| mScarlet-A-to-G-22 | GGATGTCCCAGGAGAAGGGCAGGGGGCCAC |
| mScarlet-A-to-G-24 | GATGTCCCAGGAGAAGGGCAGGGGGCCACCC |
| mScarlet-A-to-G-26 | GTCCCAGGAGAAGGGCAGGGGGCCACCCTT |
| mScarlet-A-to-G-28 | GCCCAGGAGAAGGGCAGGGGGCCACCCTTGG |
| mScarlet-A-to-G-30 | GCAGGAGAAGGGCAGGGGGCCACCCTTGGTC |
| mScarlet-C-to-U-20 | CTGCCTGTCCCACATCAATGGAGATCCAAAC |
| mScarlet-C-to-U-22 | GTGGATCTCCATTGATGTGGGACAGGCAGAT |
| mScarlet-C-to-U-24 | GATCTCCATTGATGTGGGACAGGCAGATCA |
| mScarlet-C-to-U-26 | GTCTCCATTGATGTGGGACAGGCAGATCAGG |
| mScarlet-C-to-U-28 | GTCCATTGATGTGGGACAGGCAGATCAGGGC |
| APC | GCCACTCCCAACAGGTTTCACAGTAAGCGC |
| MECP2 | GTCCGTGTCCAGCCTTCAGGCAGGGTGGGGT |
| SMN1 | GCTTCTGACCAAATGGCAGAACATTTGTCCC |
| CFTR | GCTTTCCTCCACTGTTGCAAAGTTATTGAA |

| | |
|---|---|
| HBB | gCTCTGGGTCCAAGGGTAGACCACCAGCAGC |
| SOS1 | gCATCTGTCCTTTCTACTGTATCTTCTATAT |
| MYH11 | gTGGACTGCCGCTCCTGCACCTGCGCCTCCA |
| SCN11A | gCAACAGCCCGGGTTAAGTTAATCAGGTAGA |

| RNA scaffold sequence | |
|---|---|
| 1xMS2 | CGTACACCATCAGGGTACG |
| 2xMS2 | CGTACACCATCAGGGTACGCagatGCGTACACCATCAGGGTACG |
| 1xPBSc | ttgatgta |
| 2xPBSc | ttgatgtagccttgatgta |
| 3xPBSc-Loop | ttgatgtaAGGGCCCAttgatgtaAGGGCCCAttgatgta |
| 5xPBSc-Loop | ttgatgtaAGGGCCCAttgatgtaAGGGCCCAttgatgtaAGCGCGCAttgatgtaAGCGCGCAttgatgta |
| 10xPBSc-Loop | ttgatgtaAGGGCCCAttgatgtaAGGGCCCAttgatgtaAGCGCGCAttgatgtaAGCGCGCAttgatgtaAGCCCGGAttgatgtaACCGGGCAttgatgtaAGGTACGCCAttgatgtaAGGCGTACCAttgatgtaTCGTACCCAttgatgta |
| 15xPBSc-Loop | ttgatgtaaGGCCGCTttgatgtaACGCGGTCAttgatgtaAGACCGCGAttgatgtaACTCGGAttgatgtaACCGAGATTgatgtaAGGGCCCAttgatgtaAGGGCCCAttgatgtaAGCGCGCAttgatgtaAGCGCGCAttgatgtaAGCCCGGAttgatgtaACCGGGCAttgatgtaAGGTACGCCAttgatgtaAGGCGTACCAttgatgtaTCGTACCCAttgatgta |

| | |
|---|---|
| 5xPBSc-GCC | ttgatgtagccttgatgtagccttgatgtagccttgatgtagccttgatgta |
| 15xPBSc-GCC | ttgatgtagccttgatgtagccttgatgtagccttgatgtagccttgatgtaagatttgatgtagccttgatgtagccttgatgtagccttgatgtagccttgatgtaagatttgatgtagccttgatgtagccttgatgtagccttgatgtagccttgatgta |
